# The Viral Polymerase Complex Mediates the Interaction of vRNPs with Recycling Endosomes During SeV Assembly

**DOI:** 10.1101/2020.04.23.058883

**Authors:** Emmanuelle Genoyer, Katarzyna Kulej, Chuan Tien Hung, Patricia A. Thibault, Kristopher Azarm, Toru Takimoto, Benjamin A. Garcia, Benhur Lee, Seema Lakdawala, Matthew D. Weitzman, Carolina B. López

**Author notes:** To whom correspondence should be addressed: Carolina B. López, PhD. Department of Pathobiology, School of Veterinary Medicine, University of Pennsylvania, Philadelphia, PA 19104, United States.

## Abstract

Paramyxoviruses are negative sense single-stranded RNA viruses that comprise many important human and animal pathogens, including human parainfluenza viruses. These viruses bud from the plasma membrane of infected cells after the viral ribonucleoprotein complex (vRNP) is transported from the cytoplasm to the cell membrane via Rab11a-marked recycling endosomes. The viral proteins that are critical for mediating this important initial step in viral assembly are unknown. Here we use the model paramyxovirus, murine parainfluenza virus 1, or Sendai virus (SeV), to investigate the roles of viral proteins in Rab11a-driven virion assembly. We previously reported that infection with SeV containing high levels of copy-back defective viral genomes (DVGs) generates heterogenous populations of cells. Cells enriched in full-length virus produce viral particles containing standard or defective viral genomes, while cells enriched in DVGs do not, despite high levels of defective viral genome replication. Here we take advantage of this heterogenous cell phenotype to identify proteins that mediate interaction of vRNPs with Rab11a. We examine the role of matrix protein and nucleoprotein and determine that they are not sufficient to drive interaction of vRNPs with recycling endosomes. Using a combination of mass spectrometry and comparative protein abundance and localization in DVG- and FL-high cells, we identify viral polymerase complex components L and, specifically, its cofactor C proteins as interactors with Rab11a. We find that accumulation of these proteins within the cell is the defining feature that differentiates cells that proceed to viral egress from cells which remain in replication phases.

**IMPORTANCE:** Paramyxoviruses are a family of viruses that include a number of pathogens with significant burdens on human health. Particularly, human parainfluenza viruses are an important cause of pneumonia and bronchiolitis in children for which there are no vaccines or direct acting antivirals. These cytoplasmic replicating viruses bud from the plasma membrane and coopt cellular endosomal recycling pathways to traffic viral ribonucleoprotein complexes from the cytoplasm to the membrane of infected cells. The viral proteins required for viral engagement with the recycling endosome pathway are still not known. Here we use the model paramyxovirus Sendai virus, or murine parainfluenza virus 1, to investigate the role of viral proteins in this initial step of viral assembly. We find that viral polymerase components large protein L and accessory C proteins are necessary for engagement with recycling endosomes. These findings are important in identifying viral proteins as potential targets for development of antivirals.

## INTRODUCTION

Paramyxoviruses are single-stranded negative sense RNA viruses that include viruses of clinical and global health significance, such as human parainfluenza viruses (HPIV). HPIV1 and 2 are the leading cause of croup in young children and HPIV3 is associated with bronchiolitis, bronchitis, and pneumonia (1). These infections have also been associated with the development or exacerbation of asthma and chronic airway disorders (2). Additionally, HPIVs are the etiological agent of 7% of hospitalized pneumonia cases in children in the U.S. (3), as well as in Africa and Asia (4). Although HPIVs pose a significant health burden, there are no direct acting antivirals or preventative vaccines. In order to identify targets for antiviral development, there is a need to better understand fundamental aspects of paramyxovirus biology, including mechanisms of virion assembly and particle production.

Sendai virus (SeV), or murine parainfluenza virus 1, is closely related to human parainfluenza viruses 1 and 3 with 69% homology at the RNA level between SeV and HPIV1. SeV has long served as a well-established model for understanding paramyxovirus replication. SeV genome replication occurs in the cytoplasm of infected cells (5) and virions bud from the plasma membrane. The viral genomes are tightly coated in the viral nucleocapsid protein (NP) with one NP molecule for every six nucleotides of genome (6). This viral RNA-protein complex is referred to as the viral ribonucleoprotein (vRNP) (7, 8). Viral genomes are replicated by the RNA-dependent RNA polymerase (RdRP) large protein or L(9), along with the polymerase cofactor phosphoprotein (P) which is required for recruitment of NP to nascent RNA (10). Additionally, many paramyxoviruses encode a family of accessory proteins known as C proteins (C’, C, Y1 and Y2) generated by alternative translation of the P mRNA. The C proteins were first described as interferon antagonists via binding to STAT1 (11), but have also been implicated as polymerase cofactors that regulate genome polarity and mRNA transcription (12).

The viral structural proteins include surface proteins hemagglutinin (HN) and fusion (F), which act as an attachment factor and fusion machinery respectively (13). The matrix protein (M) lines the inner membrane of the virion and is important for bridging the interactions between HN, F and vRNP complexes, and this interaction drives membrane curvature which results in the budding of virions (14, 15). In order for virion release to occur at the membrane, cytoplasmic viral genome and structural proteins must be transported to the cell surface. While F and HN are trafficked and processed through the endosomal network to the plasma membrane (16), vRNPs require transport through the cytoplasm.

Paramyxoviruses, as well as orthomyxoviruses and hantaviruses, use the recycling endosome pathway as a controlled mechanism of egress to traffic vRNPs from the infected cell cytoplasm to the cell membrane (17). Rab11a, a host GTPase associated with recycling endosomes, catalyzes bidirectional transport of endosomes along microtubule networks between the perinuclear endocytic recycling complex and the plasma membrane (18). Paramyxoviruses that rely on Rab11a for intracellular transport and viral egress include SeV (19), measles (20), mumps (21), and HPIV1 (19). While it is known that the vRNP interacts with Rab11a, it is not known which viral proteins are necessary to drive this critical interaction required for viral particle assembly. In fact, although multiple proteins including M and C have been investigated for their role in driving viral assembly their roles remain contested and poorly understood (14). For example, SeV M has been reported to interact with vRNPs in the cytoplasm and be required for recruitment of vRNPs to the membrane (22, 23), but has also been shown to traffic with surface proteins F and HN via the trans-Golgi network (24). Whether the M protein is critical for engagement of SeV vRNPs with Rab11a-marked endosomes is unknown. Further, in addition to the C proteins roles in innate immune antagonism and polymerase function, they have also been reported to be enhancers of particle formation (25, 26). Data suggest that C is critical in recruiting vRNPs to the membrane (25), and that C expression enhances association of vRNPs to membranes. Additionally, both C and M proteins have been shown independently to recruit Aip1/Alix, a protein involved in endosomal sorting and vesicle budding, to the plasma membrane to enhance virion formation (26, 27). However, the requirement for Aip1/Alix in paramyxovirus budding is contested, with evidence that absence of Aip1/Alix in cells does not have any effect on SeV particle formation (28). Thus, while both C and M have been heavily studied in the context of particle assembly, a clear picture of how these proteins function in early steps of assembly is lacking.

Defective viral genomes (DVGs) are truncated products generated during viral replication that are only able to be replicated in the presence of a full-length (FL) standard viral genome (29). DVGs of the copy-back type are robustly generated during infection with paramyxoviruses, including measles (30), mumps(31), Nipah (32) and SeV (33, 34). These DVGs are formed from the 5’ end of the anti-genome and contain complementary flanking regions. DVGs have been described to alter outcomes of infection by reducing standard viral replication, initiating viral persistence, and inducing innate immune responses (29, 35). In addition to playing a critical role in modulating viral infections, DVGs have long been used as a tool for dissecting basic processes of viral replication (6, 36), innate immune activation (37, 38), and particle production (39).

We previously reported that upon infection with DVG-containing virus populations, cells display a heterogenous phenotype with the development of subpopulations of DVG-high cells and full-length (FL)-high cells (40, 41). DVG-high cells contain higher levels of DVGs than full length genomes, and FL-high cells contain higher levels of full-length genomes than DVGs. Not only do these subpopulations have distinct transcriptional profiles (40), but they have different intracellular localizations of viral RNA(vRNA) (41). The vRNA in FL-high cells interacts with recycling endosomes and this leads to the production of both standard and defective viral particles. In contrast, the vRNA in DVG-high cells does not interact with recycling endosomes, and consequently these cells do not produce significant amounts of viral particles. These DVG-high cells do however undergo robust levels of vRNA replication, as evidenced by the large increase in DVG RNA by qPCR and vRNA fluorescent in situ hybridization (FISH) over time (40) (41). Here we take advantage of DVGs as a system to investigate initial steps that differentiate viral replication from viral particle production; namely how vRNPs interact with Rab11a. We describe the viral polymerase components L and C as a differentiating factors in FL-high cells that facilitate vRNP association with recycling endosomes and subsequent viral assembly.

## RESULTS

### M protein interacts with NP primarily at the cell surface and does not localize with Rab11a

In order to investigate whether the M protein is responsible for the association of vRNPs with recycling endosomes, we created a recombinant SeV with an HA-tag on the N-terminus of the M protein (SeV-M-HA) to study its localization during infection. We characterized this virus to ensure that the HA-tag did not result in a dramatically different growth curve from the parental SeV F1R strain (SeV-F1R). We found that while viral output was slightly lower at later time points in infection, virion production was largely unimpaired (**Fig. 1A**). We then examined the localization of M during infection. Consistent with the fact that M lines the inside of virions and budding occurs from the plasma membrane, we observed M at the plasma membrane of infected cells (**Fig. 1B-C**). Interestingly, single plane confocal images show little overlap of NP and M proteins (**Fig. 1B**). This virus allows us to define M protein intracellular distribution during replication and virion assembly.

**Figure 1.**
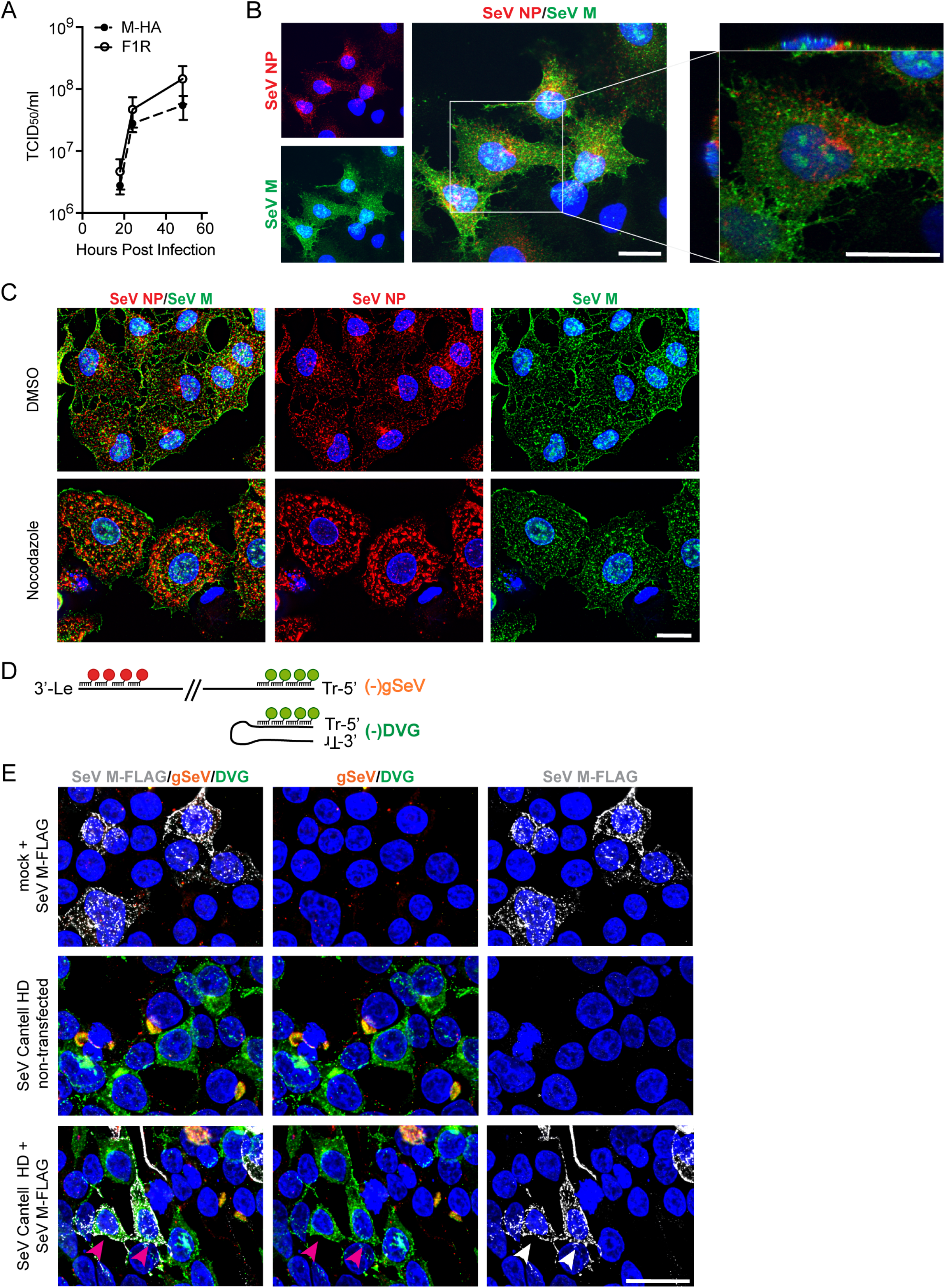
Matrix protein localization to the plasma membrane occurs independently of microtubules/Rab11a. **A)** TCID50/mL of supernatant from LLCMK2 cells infected with SeV-M-HA and SeV F1R (parental wildtype) at indicated timepoint, n = 3, graphed as mean ± SEM. **B)** A549 cells infected with SeV-M-HA for 24 hpi, immunofluorescence for SeV NP (red), HA (SeV M, green), Hoechst (nuclei, blue). Confocal 63x, 2x digital zoom. Panels 1-3 showing extended focus, panel 4 showing cropped single XY plane as well as XZ and YZ planes. **C)** A549 cells infected with SeV-M-HA, treated with nocodazole or DMSO vehicle control at 4hpi, immunofluorescence at 24 hpi for SeV NP (red), HA (SeV M, green), and Hoechst (nuclei, blue). 63x widefield deconvoluted, max projection shown. **D)** Schematic of vRNA FISH to detect (-)gSeV and (-)DVG RNA, indicating red and green probe binding regions. **E)** 293T cells infected with SeV Cantell HD then transfected with SeV M-FLAG at 6hpi, vRNA FISH with immunofluorescence for FLAG (SeV M, gray) at 24hpi. 63x widefield deconvoluted, max projection shown. Magenta arrows indicate infected and transfected cells. All images are representative of 3 independent experiments, scale bar = 20μm.

As we previously reported, when Rab11a is knocked down by siRNA or when microtubule polymerization is disrupted, the perinuclear localization of viral RNA is altered (41). To ask if M interacted with the Rab11a/microtubule pathway, we assayed localization of M upon treatment with nocodazole, a drug that prevents microtubule polymerization. In agreement with previously published data, nocodazole treatment of FL-high cells disrupted perinuclear clustering of the viral NP, indicating that vRNPs are tethered to microtubules via recycling endosomes (41). In contrast, M protein distribution was not drastically altered when cells were treated with nocodazole and it still localized at the membrane (**Fig. 1C**). These data support a model whereby the M protein is trafficked to the cell membrane independently of the microtubule network, implying that the M protein is unlikely to be critical in driving interaction between vRNPs and recycling endosomes.

It has also been reported that DVGs lead to increased degradation and turnover of M (42). Therefore, if M is the protein responsible for tethering vRNPs to recycling endosomes, it is possible that DVGs in DVG-high cells fail to interact with Rab11a due to insufficient levels of M to drive this interaction. To address this possibility, we overexpressed M-FLAG in cells infected with SeV Cantell strain with a high level of DVGs (HD) which generates a heterogenous population of DVG-high and FL-high cells (40, 41) and asked whether high levels of M were sufficient to drive a perinuclear localization of DVGs. For these experiments we used 293T cells because they allow for infection and subsequent transfection of the same cell. We used vRNA FISH to distinguish DVG from gSeV RNA (**Fig. 1D**), and confirmed that the distinct intracellular distribution of FL-high and DVG-high cells was conserved in these cells. Overexpressed M localized to the membrane of infected cells, as expected. However, overexpression of M in DVG-high cells failed to recruit DVGs to the perinuclear region with DVGs remaining distributed throughout the host cell cytoplasm (**Fig. 1E**). These data indicate that M is not sufficient to drive association between vRNPs and recycling endosomes.

Finally, to confirm that M was not responsible for driving interactions between vRNPs and recycling endosomes, we infected A549 Rab11a-mcherry cells with SeV low DVG (LD) viruses, in which all vRNPs should interact with Rab11a (41). We performed immunofluorescence for various viral proteins (**Fig. 2A**) and compared their colocalization with Rab11a (**Fig. 2B**). As previously reported (41), in SeV LD infections NP and Rab11a colocalize in the perinuclear region. In contrast, M largely does not colocalize with Rab11a (**Fig. 2B**). We also examined colocalization of Rab11a with other viral proteins, including C proteins and the polymerase L, using the previously characterized SeV-LeGFP virus (43) to visualize L protein during infection. We found that C also colocalizes with Rab11a to the same extent as NP, and that L colocalizes to an even higher degree than NP or C (**Fig. 2B**). These data indicate that at this timepoint of infection, most or all of the L protein is associated with recycling endosomes, while excess NP and the pool of C proteins which antagonize innate immunity do not. Overall, these data support a model whereby vRNPs associate with Rab11a independent of M and likely through members of the polymerase complex L and C.

**Figure 2.**
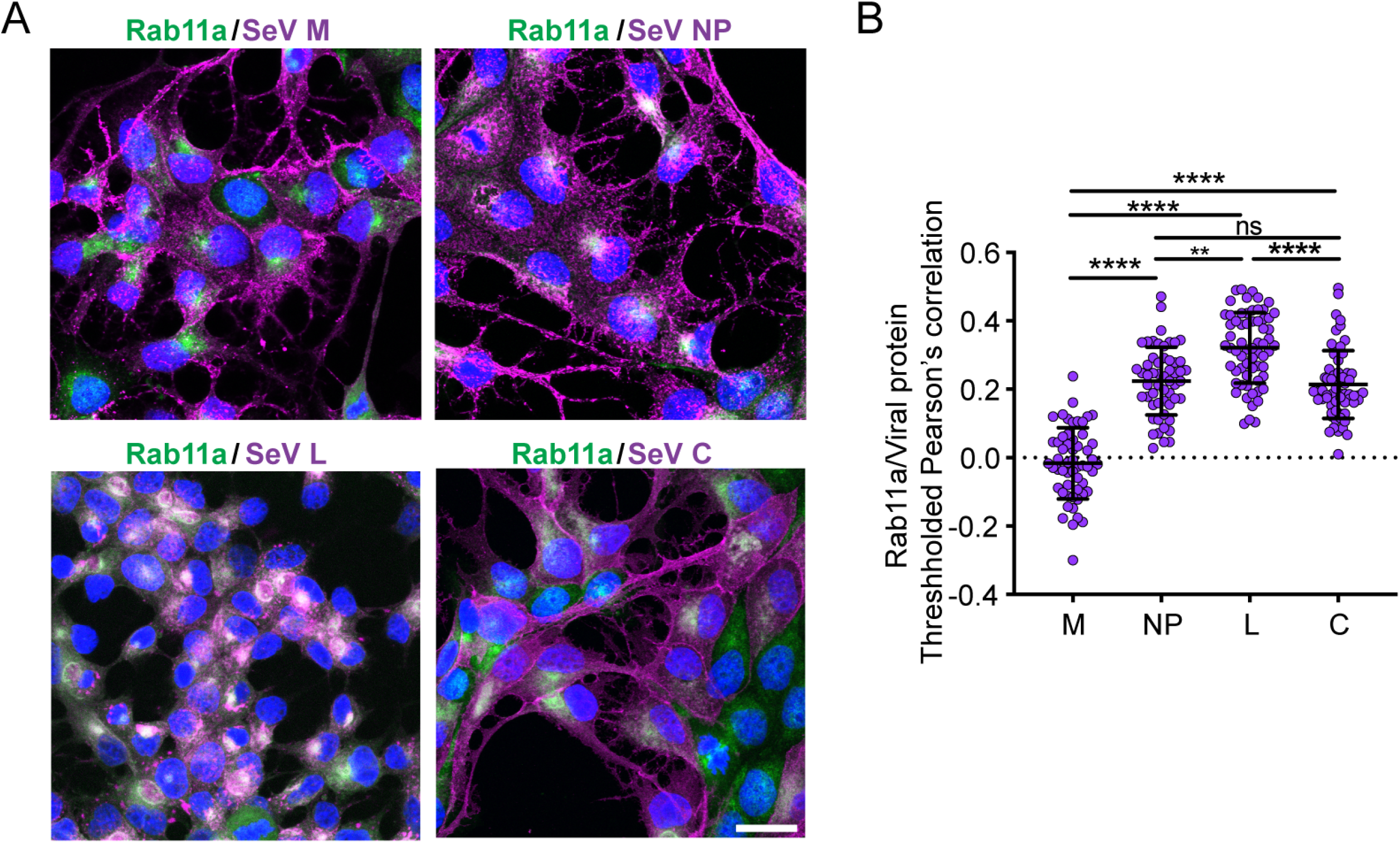
vRNP components colocalize with Rab11a but SeV M does not. **A)** A549-Rab11a-mcherry cells infected with SeV M-HA (M and NP), SeV Cantell LD (C), or SeV LeGFP (L) for 24 hours. Immunofluorescence for viral antigens (purple), and mcherry-Rab11a shown in (green). Confocal 63x images, 1.5x zoom, max projection shown. All images are representative of 3 independent experiments, scale bar = 20μm. **B)** Quantification of colocalization between viral proteins and Rab11a, pooled from three independent experiments with >20 cells analyzed per experiment. Individual cells are plotted with line at the mean and error bars represent SD. ****= p<0.0001, ** = p<0.01 by One-way ANOVA with Sidak’s multiple comparisons test.

### Nucleoprotein coverage of vRNPs is not sufficient to drive interaction with Rab11a

Since we observed that NP, L, and C proteins all colocalized with Rab11a during infection, we next sought to address which of these proteins is important for the interaction of vRNPs with recycling endosomes. First, we asked whether NP is sufficient for driving this interaction. As the most abundant protein on the vRNP, it is possible that NP coverage of the viral genome dictates engagement with Rab11a. We reasoned that if NP coverage of vRNPs was critical for interaction with Rab11a, the lack of engagement of vRNPs with Rab11a in DVG-high cells may be explained by high levels of free or partially coated vRNA. In order to investigate whether viral RNA in DVG-high cells was coated with NP, we tested for colocalization of these NP and viral genomes using RNA FISH combined with immunofluorescence for NP (**Fig. 3A**). To confirm the association of NP with DVG RNA, we quantified colocalization of 5’(-)SeV RNA with SeV NP (**Fig. 3D-F**). As shown in (**Fig. 1D**) the 5’(-)SeV probe set binds to all species of negative sense vRNA in cells. In DVG-high cells (**Fig. 3B**), the majority of vRNA recognized by 5’(-) SeV probe RNA corresponds to DVGs, but in FL-high cells (**Fig. 3C**) it corresponds primarily to FL-genome RNA as indicated. We found that there was no significant difference in colocalization of viral RNA and SeV NP between DVG-high and FL-high cells (**Fig. 3D**). Mander’s overlap coefficient (MOC) indicates the portion of a signal that overlaps with the other signal in question, regardless of signal intensity. The MOC for NP with viral RNA showed that nearly all NP in DVG-high cells overlapped with viral genomes, but a lower fraction of SeV NP overlapped with vRNA in FL-high cells, presumably due to excess free NP that is not coating viral genomes (**Fig. 3E**). The MOC for 5’(-)SeV RNA overlap with NP was equivalent between DVG-high and FL-high cells, indicating that equal proportions of viral RNA colocalized with NP (**Fig. 3F**).

**Figure 3.**
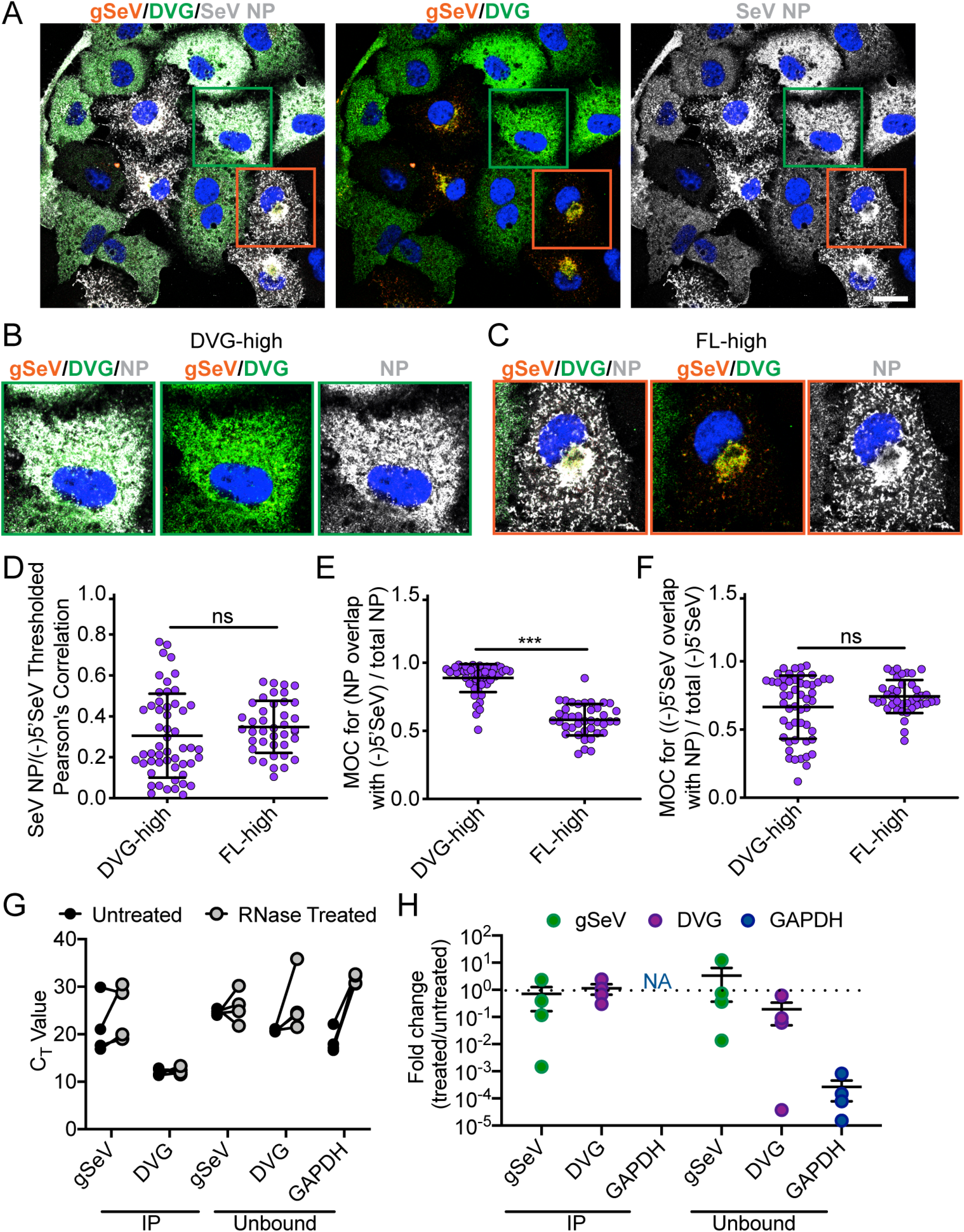
Nucleoprotein association is similar between full-length and defective viral genomes. **A)** A549 cells infected with SeV Cantell HD and then subjected to RNA FISH with immunofluorescence for NP (gray), and Hoechst staining (nuclei, blue) at 24 hpi. Confocal 63x image, 1.5x zoom, single plane image shown. Images are representative of three independent experiments, scale bar = 20μm. Green box indicates representative DVG-high cell and orange box indicates representative FL-high box. **B**) Representative cropped DVG-high cell **C**) Representative cropped FL-high cell **D)** Costes’s Pearson’s correlation of colocalization between (-)5’SeV and SeV NP in individual DVG-high and FL-high cells. **E)** Mander’s overlap coefficient (MOC) of NP with 5’ SeV in DVG-high and FL-high cells **F)** Mander’s overlap coefficient (MOC) of 5’SeV with NP in DVG-high and FL-high cells. Individual cells pooled from three independent experiments are plotted with line at the mean and error bars representing SD. **** = p<0.0001 by Mann-Whitney U-test. **G)** C_T_ values for qPCR for viral RNA, DVG RNA, and GAPDH with and without RNase A, V1, and T1 treatment after immunoprecipitation with anti-SeV NP. H) Fold change in RNA levels in RNase treated immunoprecipitation products relative to untreated. Data from four independent experiment are shown, with line at the mean and error bars representing SEM.

Though DVGs on the whole appear to be associated with NP, we wondered whether NP coating of DVGs was complete. We reasoned that there may be regions of exposed RNA that may disrupt the helical structure of the vRNP and therefore prevent interactions with host factors. To assess the integrity of the vRNP we performed an immunoprecipitation (IP) of NP from infected cells and subjected the IP product to RNase digestion with a combination of RNase A, V, and T1, targeting both single and double stranded RNA. RNA was then extracted and analyzed by RT-qPCR to compare levels of viral RNA to untreated controls. Reverse transcription was primed distal to qPCR primer sites and close to the 5’end of the genome to capture degradation at any site across the vRNA. There was no significant difference in RNA quantities between DVG RNA and full-length vRNA in treated and untreated conditions (**Fig. 3G**). We also tested unbound fractions of the pulldown, reasoning that if there was any free RNA it would not be precipitated by anti-NP antibody. Both vRNA and DVG RNA were found in the unbound fraction indicating that the IP did not precipitate all of the vRNPs. The RNA in this fraction was also not significantly different between treated and untreated conditions, particularly in comparison to an abundant cellular mRNA, GAPDH (**Fig. 3H**). These data do not preclude that there is some degree of uncoated RNA in DVG-high cells, but it indicates that the vast majority of vRNPs present during high DVG infections are sufficiently coated in NP. Our results indicate that while NP and the vRNP associate with Rab11a in FL-high cells, NP coating of viral genomes is not sufficient to drive association with recycling endosomes and is likely not providing a direct link between viral genomes and Rab11a.

### DVG driven interference leads to strong decrease in L transcripts and protein

Accumulation of DVGs interferes with standard vRNA replication and leads to a decrease in viral protein accumulation. This interference is due to DVGs competing for viral polymerase and other proteins necessary for vRNA replication (44, 45). In order to take a more systematic approach to examine of the role of viral proteins in viral assembly, we investigated whether the interference effect resulting from high DVG levels has an equal impact across viral proteins or whether certain viral mRNAs and proteins are more significantly impacted by interference. We hypothesized that proteins most interfered with by DVGs are essential for the localization of vRNPs to recycling endosomes, and therefore in their absence vRNPs mislocalize within the cytoplasm.

The transcription of paramyxovirus mRNA is directed by the RdRP beginning at the 3’ end of the genome and progressively losing processivity to generate a gradient of mRNA, with highest levels of the 3’ proximal genes and lowest levels of the 5’ genes (46, 47) produced (**Fig. 4A**). To determine whether interference with mRNA transcription by DVGs had a more drastic effect on some viral mRNAs compared to others, we performed qPCR on viral RNA from SeV LD and HD infected cell populations at 12 hours post-infection (hpi), the time at which there is the highest rate of mRNA accumulation compared to genome accumulation (12, 41). As expected, we observed a gradient of transcription with higher levels of the 3’ proximal transcripts and lowest amounts of L mRNA. Additionally, SeV HD infections had lower levels of viral mRNA levels across all genes (**Fig. 4B**). However, when comparing the levels of each mRNA across infections to normalize for differences in mRNA levels, it became clear that the levels of L mRNA were most significantly decreased (**Fig. 4C**). Because the SeV HD infections includes both FL-high and DVG-high cells, it is likely that levels of mRNA in DVG-high cells were even further reduced compared to levels of viral mRNA in LD infections, which contain mostly FL-high cells.

**Figure 4.**
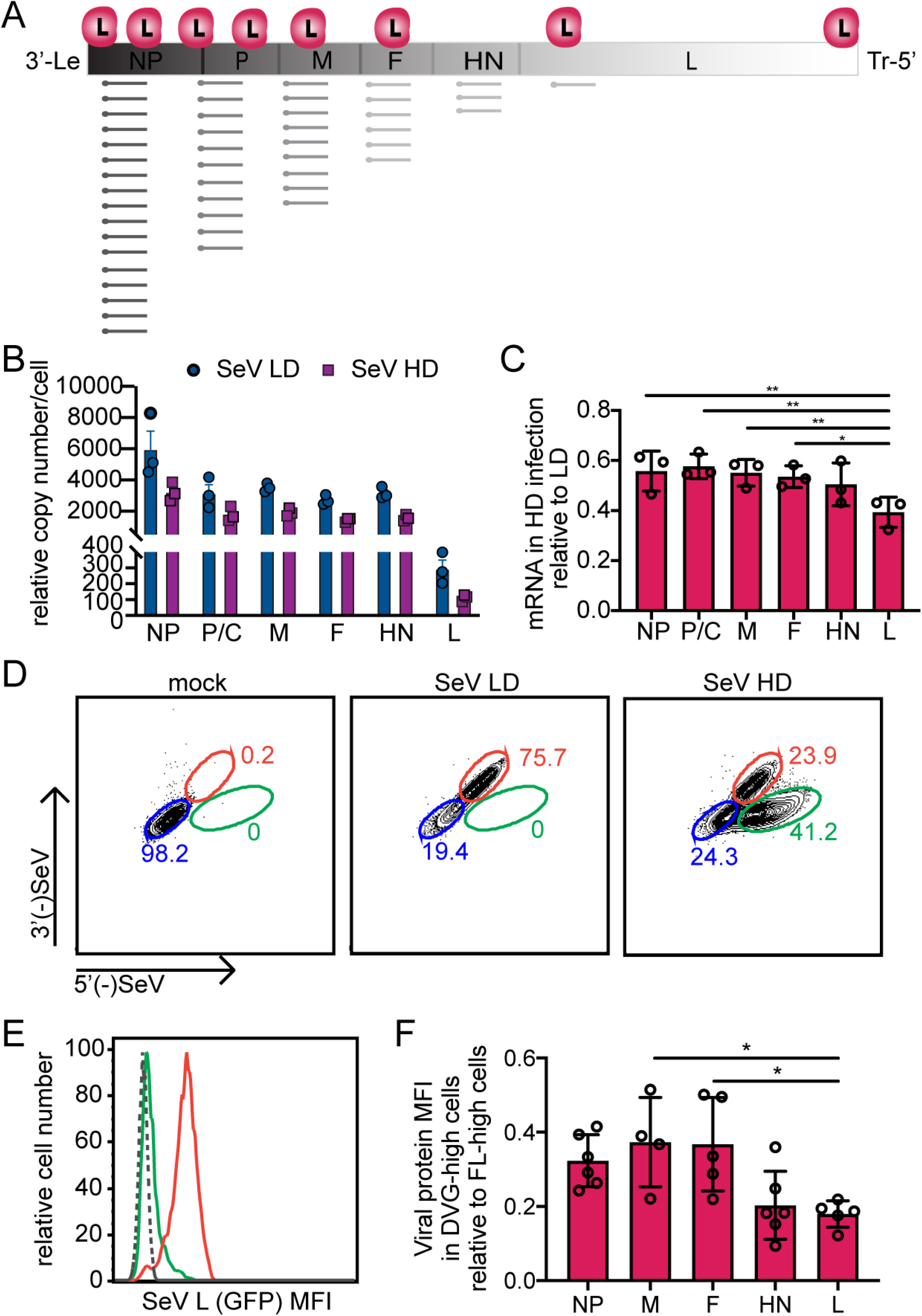
L mRNA and protein are significantly reduced in DVG-high cells compared to other viral proteins. **A)** Schematic of gene organization and gradient mRNA transcription in SeV. **(B)** A549 cells infected with SeV Cantell LD and SeV Cantell HD for 12 hours, qPCR for viral genes shown as relative copy number compared to GAPDH and **(C)** relative amount in SeV HD infections compared to SeV LD infection for each viral gene. ** = p< 0.01, * = p<0.05 by One-way ANOVA with Sidak’s multiple comparisons test. **(D-F)** A549 cells infected with SeV Cantell LD and SeV Cantell HD were subjected to RNA FISH combined with antibody staining for flow cytometery at 24 hpi. **(D)** Representative flow plots for each condition with gates for DVG-high cells highlighted in green, and FL-high cells highlighted in red, and cells below limit of detection for viral RNA by flow in blue. **(E)** Representative MFI plot showing MFI of viral proteins in different cell populations in HD infected cells with FL-high cells shown in red and DVG-high cells shown in green, mock cells shown with dashed line. **(F)** relative MFI in DVG-high cells compared to FL-high cells in SeV HD infected conditions, four independent experiments are shown with bars representing the mean and SD. NP, F, and HN were detected with monoclonal antibodies during SeV Cantell HD infection. M was detected by HA during SeV-M-HA infection and L was detected by GFP during SeV LeGFP infection in the presence of purified defective particles. * = p<0.05 by one-way ANOVA with Sidak’s multiple comparison’s test.

To determine whether decreases in viral mRNA transcription translate to lower levels of viral protein, we quantified protein levels in DVG-high and FL-high cells. SeV HD infections were subjected to RNA FISH flow coupled with immunostaining for different viral proteins. RNA FISH-flow cytometry allows for identification of discrete cell populations based on intensity of 3’- and 5’- SeV probes in each cell, using the same strategy as in imaging (**Fig. 1D**). Using this method we are able to define DVG-high and FL-high populations within the same infection (**Fig. 4D**) as validated in (40). We also assayed levels of viral protein in DVG-high and FL-high populations (**Fig. 4E-F**). Levels of protein in DVG-high cells were normalized to FL-high cell levels (**Fig. 4F**). Levels of all viral proteins were lower in DVG-high cells compared to FL-high cells, conforming with the mRNA data. These data show that the proteins that are the least abundant are most sensitive to DVG mediated interference, since there is a much greater reduction of L protein in DVG-high cells than other viral proteins (**Fig. 4F**). This observation suggests that the L protein may be an important factor in regulating interactions of vRNPs with recycling endosomes, as this is the most strongly differentiating factor between DVG-high and FL-high cells.

### SeV C proteins interact with Rab11a

We next used an unbiased approach to determine which viral proteins interact with Rab11a. To do this, we infected Rab11a-GFP A549 cells with SeV LD followed by immunoprecipitation (IP) of Rab11a using antibodies against GFP at 12 and 24 hpi to identify interacting proteins (**Supplementary Table 1**). By using SeV LD virus stocks we ensure that all cells are have vRNPs that are interacting with Rab11a. We also performed mass spectrometry on whole cell lysates in order to obtain the total quantification of all viral protein levels (**Supplementary Table 2**). We identified viral proteins in the whole cell lysate at 12 hpi, with NP showing the highest abundance, and L and accessory proteins C and V at the lowest abundance (**Fig. 5A**). This is consistent with the abundance of viral mRNA (**Fig. 4B**). In the IP of Rab11a, we identified high levels of NP, M, and C proteins (**Fig. 5B**). To define which proteins are specifically enriched by the Rab11a IP versus those just increasing in abundance after viral infection, we normalized the abundance of each protein identified in IP by their abundance in the whole cell lysate. C proteins were the most highly enriched viral proteins in IP after normalization **(Fig. 5C)**. Similar results are observed at 24 hpi, where all viral proteins were more abundant but with similar ranks as 12 hpi (**Fig. 5D-F**). We also compared the log2 fold change of proteins identified by IP at12 hpi (**Fig. 5G**) and 24 hpi (**Fig. 5H**) and identified the C proteins as the most significantly enriched viral protein interacting with Rab11a compared to mock at both time points. Additionally, L protein was identified as significantly enriched at 24 hpi. To validate these results, we performed western blot analysis and probed for Rab11a and various viral proteins (**Fig. 5I**). As expected, Rab11a was enriched after IP, while the viral protein P was not seen, corresponding with its absence in the mass spectrometry analysis. C protein had the highest ratio of IP to input, strongly suggesting that the C protein most directly interacts with Rab11a. Both NP and M proteins were present in the IP at moderate levels, which likely reflects high levels of NP on the vRNP which may be co-precipitated and potential interactions between Rab11a and M which may occur at the plasma membrane in late stages of particle formation (see Figure 1). Since SeV generates four C proteins that are described to have discrete functions (48), we asked whether we could identify which of the C proteins was interacting with Rab11a. The four C proteins range in size from 175 to 215 amino acids and they share a common 175 amino acid C-terminus. We mapped the peptide reads identified by mass spectrometry in the IP to the C proteins (**Fig. 5J**) and found they were all from the shared 175 amino acid region. Therefore, we could not conclude which specific protein(s) were identified. Overall, the identification of C proteins as a likely interactor of Rab11a by immunoprecipitation and subsequent mass spectrometry supports results showing high degrees of colocalization as shown in (**Fig. 2A-B**) and suggests C proteins play an important role in directing interaction of vRNPs with recycling endosomes.

**Figure 5.**
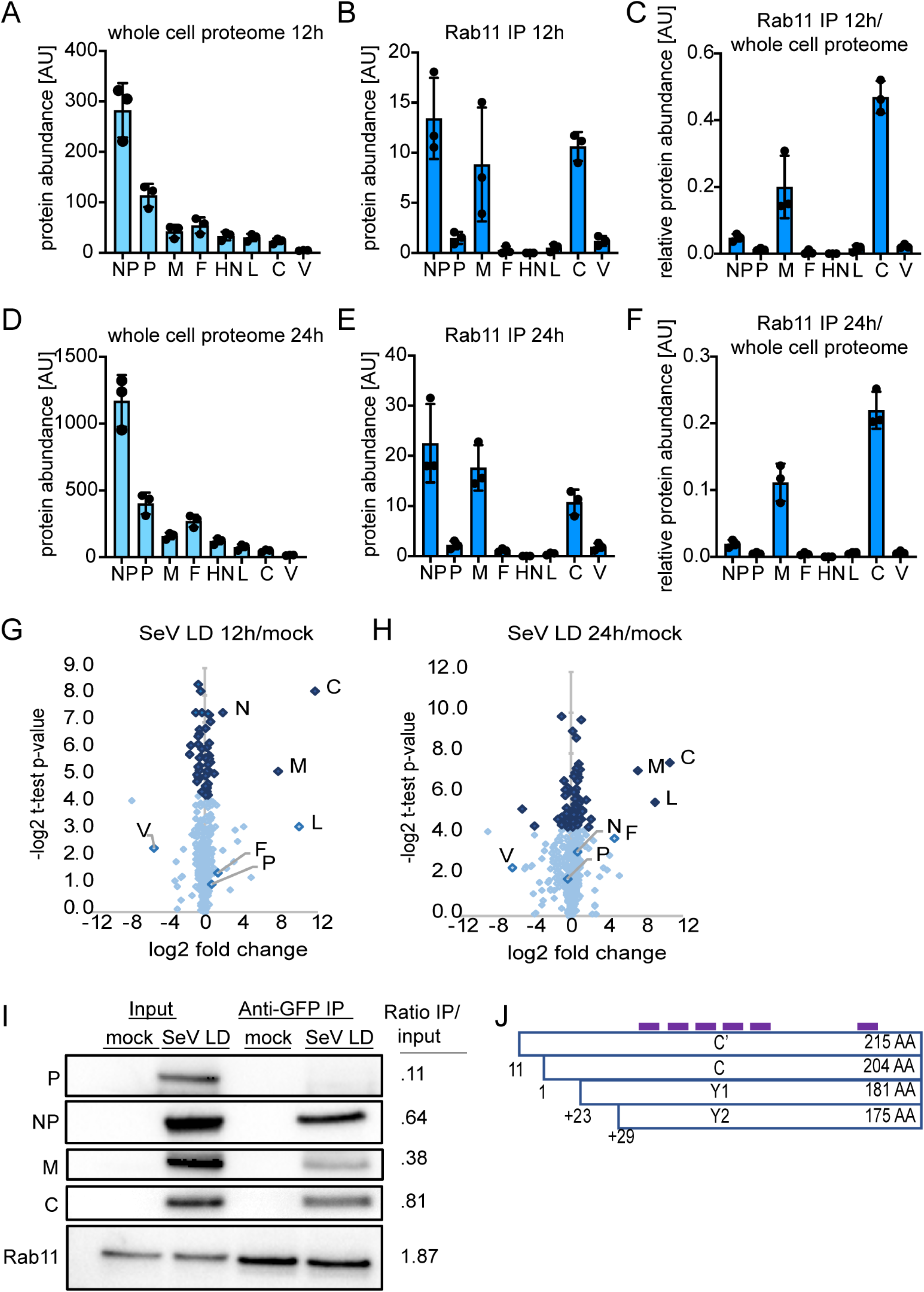
C proteins identified to interact with Rab11a by immunoprecipitation and mass spectrometry. **A)** Viral protein abundance identified by mass spectrometry in the whole cell lysate at 12 hpi. **B)** Viral protein abundance identified by IP of Rab11a-GFP at 12 hpi. **C)** Relative ratio of IP/whole cell lysate of viral proteins at 12 hpi. **D)** Viral protein abundance identified by mass spectrometry in the whole cell lysate at 24 hpi. **E)** Viral protein abundance identified by IP of Rab11a-GFP at 24 hpi. **F)** Relative ratio of IP/whole cell lysate of viral proteins at 24 hpi. A-F, three independent experiments are plotted with bar representing mean, error bar representing ±SD. Volcano plot highlighting proteins enriched in Rab11a-GFP IP at 12 hpi **(G)** and at 24 hpi **(H)** compared to mock. Abundance of each protein identified in IP was normalized by their abundance identified in the whole cell lysate. In order to estimate an enrichment of proteins detected only in mock or infection, missing values were imputed. Data are pooled from three independent experiments. Viral proteins are indicated. Dark blue dots represent p<0.05 and light blue dots represent p>0.05 by student’s t-test. (**I**) Validation of mass spectrometry by western blot probing for Rab11a or viral proteins at 24 hpi and quantified for relative band intensity of IP over input. (**J**) Schematic of C proteins with locations of peptides identified by mass spectrometry in IP samples marked in purple.

### Occupancy of polymerase proteins on viral genomes is a differentiating factor for vRNPs interaction with Rab11a

Because we found that the L protein was lowest in DVG-high cells when comparing DVG-high and FL-high cells and identified its cofactor C by mass spectrometry as a Rab11a interacting partner, we considered whether differences in polymerase component levels may drive differences in engagement with Rab11a in DVG-high and FL-high cells. To ask whether polymerase was associated with viral RNA in DVG-high cells we performed RNA FISH combined with immunofluorescence. To visualize L we used a recombinant SeV with the L protein C-terminally fused to eGFP (SeV LeGFP) that has been previously described (43) and the addition of purified defective particles containing DVG-546, the dominant DVG in SeV Cantell. Previous studies have validated the ability of other strains of SeV to replicate DVGs from heterologous strains (41). During infection with SeV LeGFP in the presence of purified defective particles (pDPs), we see the establishment of DVG-high cells, and strikingly these cells have very low levels of L protein (**Fig. 6A)**, recapitulating flow cytometry results that showed large differences in the amount of L protein in DVG-high (**Fig. 6B**) and FL-high cells (**Fig. 6C**). Additionally, it appears that a majority of DVG RNA in DVG-high cells is not bound by L protein. Indeed, colocalization of 5’(-)SeV RNA and L was significantly lower in DVG-high cells compare to FL-high cells (**Fig. 6D**). The MOC of L is equivalent in DVG-high and FL-high cells, indicating that the same proportion of L in each cell is interacting with vRNA. The mean score nearing 1 indicates that the small amount of polymerase present in those cells is colocalized with DVG RNA (**Fig. 6E**). However, the MOC of 5’ (-) SeV RNA with SeV L was significantly lower in DVG-high cells compared to FL-high cells, indicating that a smaller fraction of viral RNA overlaps with L in DVG-high cells than in FL-high cells and that there are many DVG RNAs that do not colocalize with detectable polymerase in DVG-high cells (**Fig. 6F**). The small amount of L protein in DVG-high cells is likely associated with DVGs for replication, but the majority of DVGs within the cytoplasm of DVG-high cells are not associated with detectable levels of L. This strong difference in L interaction with viral RNA in DVG-high and FL-high cells indicates that L protein may be critical in driving vRNPs to interact with Rab11a.

**Figure 6.**
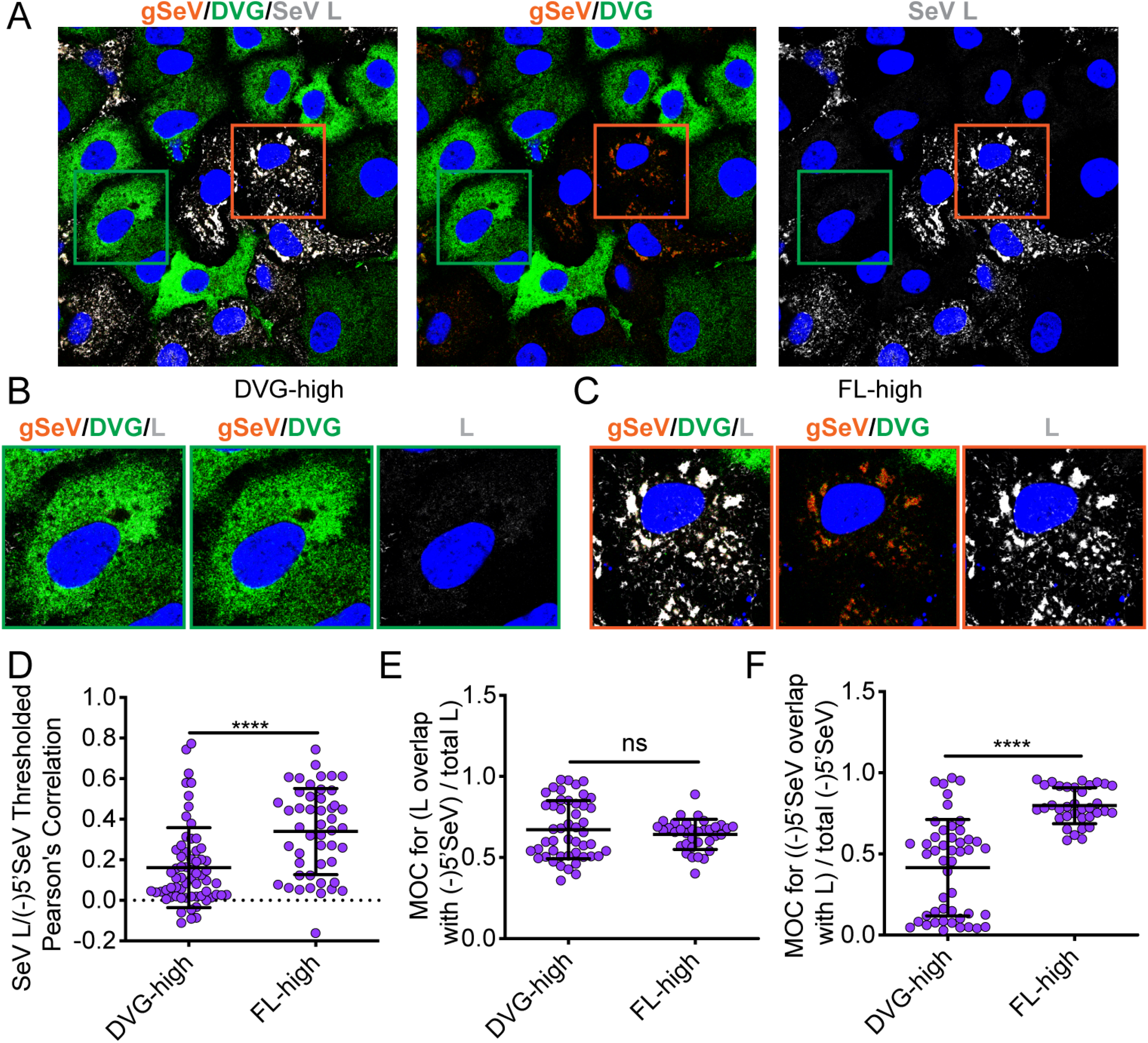
L protein interaction with vRNP distinguishes between DVG-high and FL-high cells. **A)** A549 cells infected with SeV LeGFP supplemented with purified DPs (HAU 20) for 24 hours and then subjected to RNA FISH with immunofluorescence for GFP (L, gray), and Hoechst staining (nuclei, blue). Confocal 63x image, 1.5x zoom, single plane image shown. Images are representative of four independent experiments, scale bar = 20μm. Green box indicates representative DVG-high cell and orange box indicates representative FL-high box. **B**) Representative cropped DVG-high cell. **C**) Representative cropped FL-high cell. **D)** Costes’s Pearson’s correlation of colocalization between (-)5’SeV and SeV NP in individual DVG-high and FL-high cells. **E)** Mander’s overlap coefficient of L (GFP) with 5’ SeV in DVG-high and FL-high cells **F)** Mander’s overlap coefficient of 5’SeV with L (GFP) in DVG-high and FL-high cells. Individual cells pooled from three independent experiments are plotted with line at the mean and error bars representing SD, **** = p<0.0001 by Mann-Whitney U-test.

Since engagement with the polymerase seems to be a differentiating factor between DVG-high and FL-high cells, we sought to investigate whether similar differences were found with C protein as well. Using RNA FISH combined with immunofluorescence for C protein, we observed similar results to those seen with SeV L, with low levels of C protein in DVG-high cells and a perinuclear localization of C protein in FL-high cells, as well as at the plasma membrane (**Fig. 7A**). These results suggest that at later timepoints of infection, C protein associates with vRNA and presumably L. Quantification of colocalization revealed that there is a significantly greater degree of colocalization of C and 5’(-)SeV RNA in FL-high cells (**Fig. 7C**) compared to DVG-high cells (**Fig. 7B**). This observation suggests that like with L protein, many DVG vRNPs are not occupied by C protein (**Fig. 7D).** MOC of SeV C reveals that higher proportions of C are colocalized with 5’ (-) SeV RNA in DVG-high cells than in FL-high cells (**Fig. 7E**). However, MOC of 5’ (-) SeV RNA with SeV C indicates that in FL-high cells, a high proportion of vRNA overlaps with C, while in DVG-high cells, much of the vRNA does not overlap with C proteins (**Fig. 7F**). These data indicate, that similarly to L protein, low levels of C proteins in DVG-high cells seem to colocalize with DVG-RNA but that very low levels of C protein in the cell leave many RNAs unbound to free RNA. Taken together, these data suggest that lack of accumulation of C and L proteins in DVG-high cells, and therefore lack of C and L proteins bound to vRNPs precludes these vRNPs from interacting with Rab11a.

**Figure 7.**
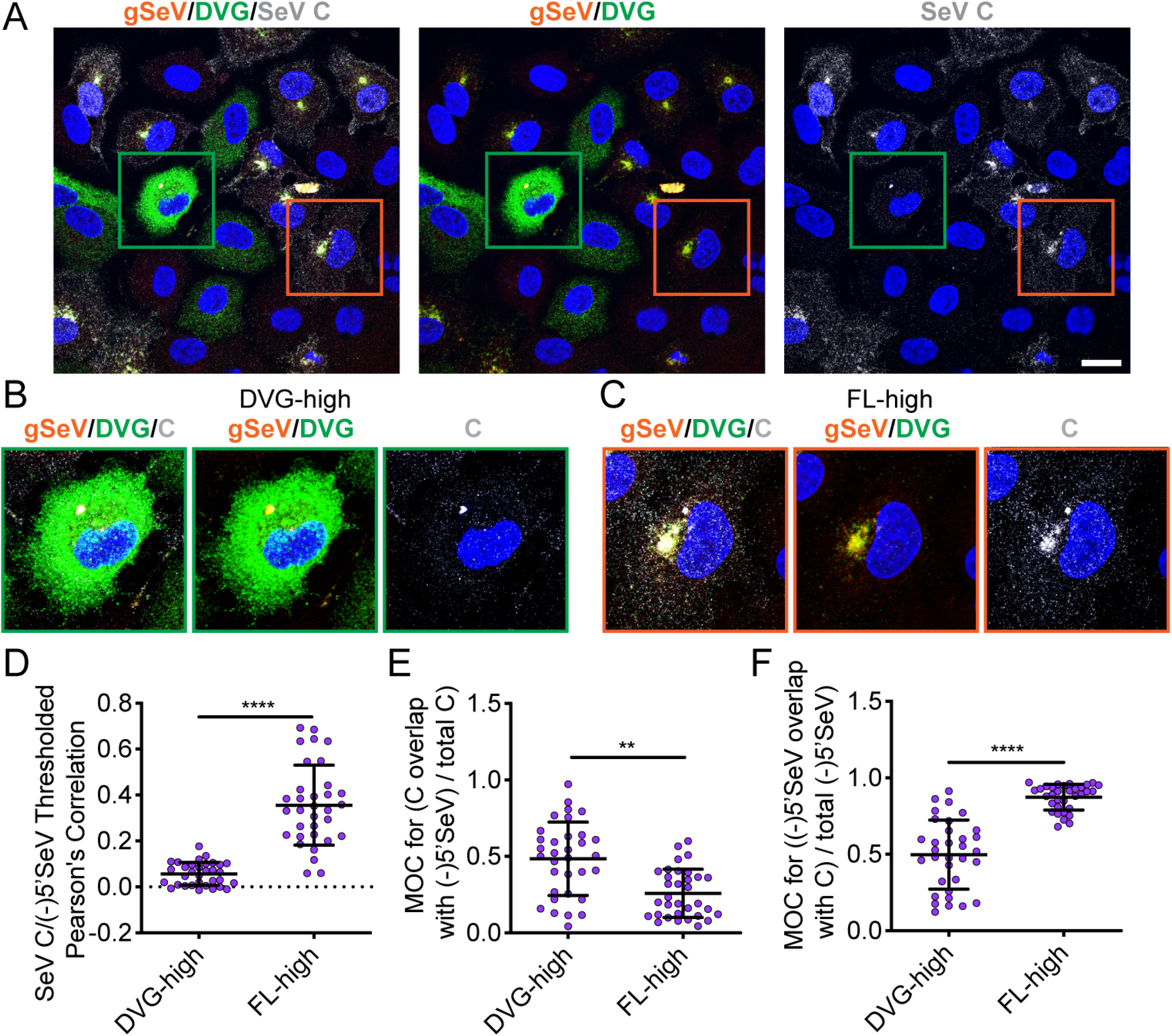
C protein interaction with vRNP distinguishes between DVG-high and FL-high cells. **A)** A549 cells infected with SeV Cantell for 24 hours and then subjected to RNA FISH with immunofluorescence for C (gray), and Hoechst staining (nuclei, blue). Confocal 63x image, 1.5x zoom, single plane image shown. Images are representative of four independent experiments, scale bar = 20μm. Green box indicates representative DVG-high cell and orange box indicates representative FL-high box. **B**) Representative cropped DVG-high cell **C**) Representative cropped FL-high cell **D)** Costes’s Pearson’s correlation of colocalization between (-)5’SeV and SeV C in individual DVG-high and FL-high cells. **F)** Mander’s overlap coefficient of C with 5’ SeV in DVG-high and FL-high cells **E)** Mander’s overlap coefficient of 5’SeV with C in DVG-high and FL-high cells. Individual cells pooled from two independent experiments are plotted with line at the mean and error bars representing SD **** = p<0.0001 by Mann-Whitney U-test.

## DISCUSSION

We propose a model of Sendai virus assembly in which polymerase components including the viral polymerase L and its cofactor C are critical for the interaction between viral RNPs and Rab11a. This interaction is a crucial initial step in viral particle production. In taking advantage of the difference in viral particle production in DVG-high and FL-high cells, we were able to parse viral components that are critical for replication from those that are critical for viral assembly. DVG-high cells are able to carry out robust levels of viral genome replication using limited amounts of L but are unable to produce virions because they do not engage with host proteins critical for this process (41). We observed that DVG interference with viral protein expression critically diminishes levels of the viral polymerase L in DVG-high cells. We propose that L/C accumulation above a certain threshold is required for interaction of the vRNP with recycling endosomes and the cellular trafficking machinery. Furthermore, using mass spectrometry we identified the C proteins as the most highly enriched viral proteins interacting with Rab11a. From these data we propose that recycling endosomes interacts with C proteins when it is associated with L, and that C-L polymerase complexes interact with vRNPs to tether them to recycling endosomes (**Fig. 8**).

**Figure 8.**
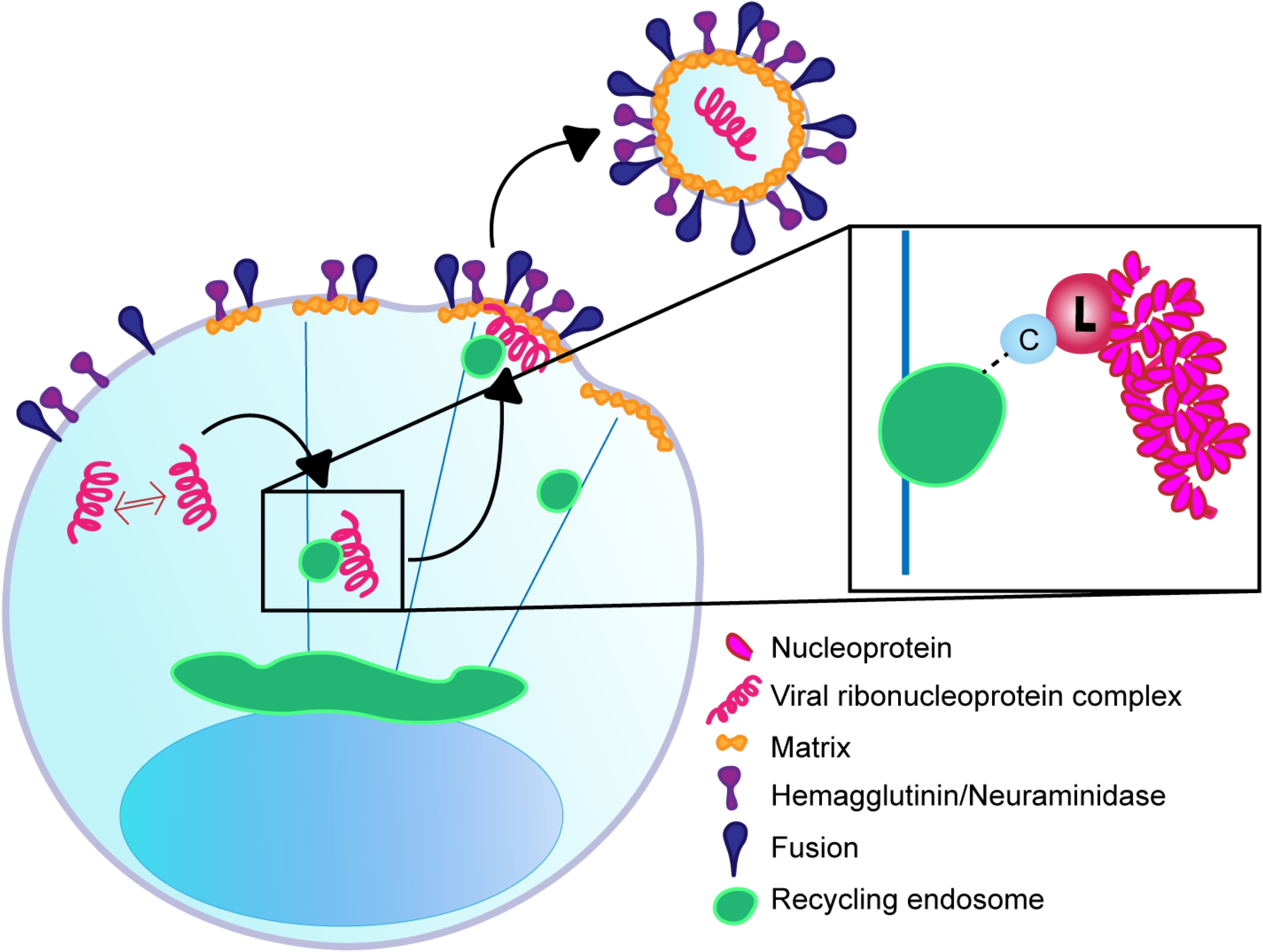
Model of SeV vRNP interaction with Rab11a. Viral replication occurs in the cytoplasm and upon accumulation of sufficient levels of L and C proteins, vRNPs interact with Rab11a-marked recycling endosomes via C and L proteins, to enable trafficking to the plasma membrane.

It remains to be determined whether paramyxoviruses interact directly with recycling endosomes via endosomal membrane interactions, with the Rab11a GTPase itself, with family-interacting proteins (FIPs) associated with Rab11a or via another mechanism. The different Rab11-FIPs direct engagement of Rab11a-marked recycling endosomes with dynein or myosin Vb motor proteins, and specific FIPs have been identified as important for budding of different viruses. For example Rab11-FIP2 is important for respiratory syncytial virus budding (49) and influenza relies on Rab11-FIP3 for filamentous particle production (50). The Rab11a-FIPs used for trafficking of SeV are unknown, and identifying the viral protein that mediates interaction with recycling endosomes will aid further study of these dynamics.

Negative-sense RNA viruses require packaging of RdRPs within virions in order to initiate infection. Indeed, SeV L proteins have been shown to associate with viral nucleocapsids isolated from virions (51). Additionally, C proteins have been found to be tightly associated with nucleocapsid structures both inside the cell as well as when isolated from SeV virions, with approximately 40 C proteins per vRNP (52). The observed role of polymerase components in assembly are consistent with what was observed in infections with the orthomyxovirus influenza, where influenza virus RNPs interact with Rab11a via one of its polymerase components PB2 (53, 54). As has been suggested for influenza, having the SeV polymerase complex initiate packaging via interactions with Rab11a could be a mechanism to ensure that only vRNPs that contain a RdRP, and are thus infectious, can be packaged into virions.

Most paramyxoviruses encode a number of accessory proteins from the P gene, either through RNA editing leading to interferon antagonist V and W families, or by translating alternate reading frames from the P mRNA (55) to generate C family members. While we propose that C proteins are critical for engagement with Rab11, we suggest that in the absence of sufficient accumulation of L protein, vRNPs would be unable to interact with Rab11. Thus, even if, for example, levels of C protein were higher in DVG-high cells, we would not expect to see C interacting with vRNPs in the absence of L. Additionally, the specific mechanisms that dictate translation of the C proteins (C, C’, Y1 and Y2) from the P mRNA are unknown. Therefore, how DVG-mediated interference affects C protein accumulation is unclear. While the C proteins have been described to have discrete functions in regulating pathogenesis and synthesis of mRNA and vRNA synthesis (48), our results are unable to distinguish whether a specific C protein is required for interaction with Rab11a or if they are interchangeable. The specific C proteins required for SeV assembly, and whether C family proteins of other paramyxoviruses play similar or distinct roles in viral assembly are as of yet unknown. Discerning their function in assembly could provide specific targets for development of direct acting antivirals.

We also investigated the role of SeV M in viral assembly. M lines the inside of virions and interacts directly with nucleoproteins (23, 56). These contacts between M and the nucleoproteins drive virus-specific viral particle production, since studies combining M and nucleoproteins from the closely related SeV and HPIV1 fail to lead to particle production (57). While M-NP contacts are important for particle production, it is unknown whether M is delivered to the cell membrane with nucleoproteins or independently, and whether its transport relies on Rab11a (14). Our experiments indicate that M traffics independently to the plasma membrane and is not the key mediator in vRNP interactions with Rab11a. Our data is supported by previous reports that indicate that M traffics with F and HN through the trans-Golgi network (TGN) to the cell membrane (24). Although we identified M in the Rab11a IP-mass spectrometry data, this result could not be validated by western blot. This may reflect interactions between M and Rab11a occurring at the plasma membrane, at the TGN or the ER. If M is trafficked to the membrane with surface proteins F and HN as proposed by (24), Rab11a may be interacting with M near the TGN. There is evidence that the rhabdovirus vesicular stomatitis virus (VSV) glycoprotein, which is processed in the TGN, requires Rab11a for proper sorting to the plasma membrane (58). There is also evidence that Rab11a is recruited to the ER during influenza virus infection to facilitate transport of vRNPs (59). Therefore, it is possible that SeV M, F, and HN may interact with Rab11a during their transit from the ER through the TGN to the plasma membrane.

It is well established that DVGs exert effects on viral infections through interfering with standard viral genomes by competing for structural proteins or viral polymerase (29). DVGs have been reported to lower levels of viral proteins globally within infected cells and for that reason have been investigated as antivirals themselves (60). Whether all viral proteins are equally affected by interference has not been previously determined. While it has been shown that infections with SeV HD leads to lower levels of M protein, these lower levels were attributed to increased degradation of M (42). Presence of DVGs have also been shown to lead to lower levels of HN on the surface of infected cells (61). We performed a systematic investigation of the effects of interference on levels of mRNA and protein levels by directly comparing FL-high cells and DVG-high cells from a heterogenous infection. Our results support previous finding of lower levels of M and HN in the presence of DVGs but reveal that DVG interference with L mRNA transcription leads to significantly lower amount of mRNA and subsequent lower protein translation in DVG-high cells. Since L is the lowest abundance mRNA and protein, DVG-driven interference appears to have the highest impact on its abundance. The gradient expression of mRNA as well as copy-back DVGs are unique to single-stranded negative sense RNA viruses. However, other RNA viruses, such as influenza produce deletion DVGs and are known to be sensitive to DVG-driven interference. It has been shown that DVGs generated from certain segments can exert different levels of interference pressure (62), but not whether interference particularly affects the accumulation of certain mRNAs or proteins. The role that DVG-mediated interference may play in restricting viral progeny production via reduction of certain key proteins in other viral families remains to be investigated.

Overall, using DVGs as a tool to understand differences in viral replication and viral assembly we have shown that polymerase cofactors may be critical in inducing primary steps of viral particle production. While we propose that C and L proteins are essential for driving contact between vRNPs and Rab11a during SeV infection, whether intricacies of viral assembly are conserved across paramyxoviruses is unknown. Many studies suggest that there are at least subtle differences amongst the viruses in their intracellular localization, transport, and eventual assembly (14). Because DVGs have been identified from most paramyxoviruses, comparisons between DVG-high and FL-high cells during infection may generally be useful in parsing proteins required for viral replication from those required to initiate virion assembly of other members of the viral family.

## MATERIALS AND METHODS

### Cells

A549 (human type II pneumocytes, ATCC CCL-185), HEK 293T/17 (ATCC^®^ CRL-11268) and LLCMK-2 (monkey kidney epithelial cells, ATTC CCL-7) were maintained in Tissue Culture Medium (DMEM (Invitrogen) supplemented with 10% Fetal Bovine Serum (Sigma), Gentamicin (ThermoFisher), Sodium Pyruvate (Invitrogen), L-glutamine (Invitrogen)) at 7% CO2, 37°C. A549 Rab11a-GFP and A549 Rab11a-mCherry cells were generated as described in (54). Cells were tested monthly for mycoplasma MycoAlert Mycoplasma detection kit (Lonza) and treated with Mycoplasma Removal Agent (MP Biomedical) upon thawing.

### Viruses

Sendai virus Cantell was grown in 10-day embryonated specific pathogen free chicken eggs (Charles River) for 40 hours before allantoic fluid was collected. Low DVG and high DVG stocks were produced as previously described (38). Defective particles (pDPs) containing DVG-546 were purified from allantoic fluid infected with SeV Cantell by density ultracentrifugation on a 5-45% sucrose gradient, using TCID_50_ to confirm the absence of virus and direct hemagglutination to detect particles(34). SeV-LeGFP is a recombinant Enders strain of SeV with eGFP fused to the C-terminus of the L protein described in (43). SeV-LeGFP cells was grown for 72 hours in LLCMK2 cells. SeV-M-HA virus was generated with an HA tag at the N-terminus of the M protein in a recombinant SeV F1R background with an eGFP reporter. Virus was rescued by transfection in to BSRT7 cells as described (63) and passaged blindly three times. Virus was then grown for 5 days in LLCMK2 cells. All viruses were titrated using 1:10 serial dilution in triplicate in LLCMK2 cells in the presence of 2μg/mL TPCK trypsin (Worthington Biomedical) for 72 hours and TCID50/mL was determined by the Reed-Meunch method based on hemagglutination of 0.5% washed chicken red blood cells (Lampire) by supernatant to indicate presence of infection.

### Plasmids and Transfection

SeV-M-FLAG plasmid encodes codon optimized SeV Matrix with a 3X N-terminal FLAG tag in the pCMV-3Tag-1A vector. Cells in antibiotic free media (8×10^5^/6-well) were transfected 6 hours post infection with 2μg plasmid and 6μl Lipofectamine 2000 (Invitrogen) per well diluted in Opti-MEM (Invitrogen).

### Viral Infections

Infections were performed at a multiplicity of infection of 1.5 TCID_50_/cell unless otherwise specified. Cells were seeded to reach 80% confluency upon day of infection (4×10^5^ A549/well in 6 well plates). Prior to infection cells were washed twice with PBS. Virus was diluted in infection media (DMEM (Invitrogen) supplemented with 35% BSA (Sigma-Aldrich), Pen/Strep (Invitrogen), 5% NaHCO_3_ (Sigma-Aldrich) and low volume inoculum was added to cells for 1 hour incubation at 37°C, with shaking every 15 minutes. Cells were then replenished with 2% FBS Tissue Culture Medium. Viral infections in the presence of pDPs were performed by adding pDPs at a dilution of 20 direct hemagglutination units/4×10^5^ cells along with virus.

### Drug Treatment

Four hours post infection, media was removed and replaced with 2% FBS tissue culture media containing 2μg/mL Nocodazole (Sigma-Aldrich) for duration of infection.

### Immunofluorescence

Cells were seeded on No. 1.5 glass coverslips (Corning) overnight prior to infection. Post infection coverslips were rinsed in PBS then fixed for 10 minutes in 4% Paraformaldehyde in PBS (Electron Microscopy Sciences) for 10 min. Cells were permeabilized with 0.2% Triton-X in PBS (Sigma-Aldrich) for 10 minutes. Primary and secondary antibodies were diluted in 3% FBS in PBS and incubated at RT for 1.5 hour and 1 hour, respectively. Nuclei were stained with Hoechst prior to mounting coverslips on slides using Fluoromount-G (ThermoFisher). Primary antibodies: mouse anti-SeV NP (clone M73/2 a kind gift from Alan Portner – directly conjugated with Dylight 594 NHS Ester (ThermoFisher)), rabbit Anti-HA-Tag (CST C29F4), mouse Anti-FLAG 9A3 (CST 8146), rabbit Anti-GFP (ab6556), mouse anti-C (clone P16, 96-6, from Toru Takimoto). Secondary antibodies: Goat anti-Rabbit IgG (H+L) Secondary Antibody, Alexa Fluor 488 (Invitrogen), Goat anti-mouse IgG (H+L) Secondary Antibody, Alexa Fluor 647 (Invitrogen); Goat anti-mouse IgG (H+L) Secondary Antibody, Alexa Fluor 488 (Invitrogen)

### RNA-FISH

Custom probe sets were designed against the SeV genome (described in(41)) and conjugated to Quasar 570 (3’(-)SeV) and Quasar 670 (5’(-)SeV) dyes (LGC Biosearch). For RNA-FISH microscopy, cells were seeded on No. 1.5 glass coverslips (Corning) prior to infection. Post-infection coverslips were rinsed with PBS then fixed in 3.7% formaldehyde (ThermoFisher) for 10 minutes. Cells were permeabilized in 70% EtOH for 1 hour at room temperature then washed in wash buffer (2X SSC (ThermoFisher), 10% formamide (ThermoFisher) in nuclease free water), and then subjected to hybridization with FISH probes. For hybridization, probes were diluted to 2.5nM in hybridization buffer (wash buffer + dextran sulfate) and applied to slides. Slides were incubated with probe overnight at 37°C in a humidified chamber. Prior to imaging, slides were washed in wash buffer twice (once with Hoechst to stain nuclei) then 2X SSC. Cells were mounted in ProLong Diamond Antifade Mountant (ThermoFisher) and cured overnight at room temperature prior to imaging. RNA FISH combined with immunofluorescence was modified by staining cells with antibody after permeabilization with EtOH. Cells are stained with antibody in 1% BSA (ThermoFisher) PBS with RNaseOUT (Invitrogen) for 45 minutes for 1° Ab and 40 minutes for 2° Ab. Cells were then post-fixed for 10 minutes in 3.7% formaldehyde prior to hybridization.

### RNA FISH Flow Cytometry

Cells were harvested post infection by trypsinization (ThermoFisher) and centrifugation. Cell pellets were washed in 1% FBS in PBS one time. Cell pellets were then resuspended in 10mL ice cold methanol (ThermoFisher) and fixed and permeabilized on ice for 10 minutes before pelleting at 1500rpm for 5 minutes. Pellets were then resuspended in 1mL wash buffer per sample and transferred to a microcentrifuge tube where they were washed twice. For hybridization probes were diluted to 25nM in 100μl hybridization buffer (wash buffer + dextran sulfate) and cell pellets were resuspended in hybridization buffers and incubated overnight at 37°C. For antibody staining, primary antibodies were added to the hybridization buffer and incubated overnight. Post-hybridization cell pellets were washed in wash buffer twice, then washed in 2X SSC before being resuspended in anti-fade buffer (2X SSC, 0.4% glucose (Sigma), Tris-HCL pH 8.0 (USB Corporation) with Catalase (Sigma) and Glucose Oxidase (Sigma)). For secondary antibody staining, antibody was added to wash buffer prior to washing with 2X SSC and incubated at 37°C for 30 minutes. Primary antibodies: mouse anti-SeV NP (clone M73/2 a kind gift from Alan Portner), mouse-anti-SeV F (clone 11H12), mouse anti-SeV HN (clone 1A6), rabbit anti-HA-Tag (CST C29F4), rabbit anti-GFP (ab6556). Secondary antibodies: Goat anti-Rabbit IgG (H+L) Secondary Antibody, Alexa Fluor 405 (Invitrogen), Goat anti-mouse IgG (H+L) Secondary Antibody, Alexa Fluor 405 (Invitrogen). Flow cytometry was performed on a BD LSRFortessa and compensation and gating was done in FlowJo_v9.9.4 (BD Biosciences). Mock and SeV LD infections were used to inform gating of SeV HD DVG-high and FL-high populations.

### Microscopy and Image Analysis

Images were acquired on a Leica SP5-II Laser Scanning Confocal with 63X (1.40-0.60 NA) and 100X (1.46 NA) oil-immersion objectives with pixel size 70×70nm for FISH immunofluorescence, with z-step size of 0.13μm, with 15 slices total and 2048×2048 image format. Immunofluorescence images (Rab11a colocalization) were acquired with the same parameters except for a pixel size of 40×40nm. Images were processed in Volocity (Perkins-Elmer) and deconvolution when performed was done with Huygen’s Essential Deconvolution Wizard using theoretical point spread function. Colocalization was quantified in Volocity over total cell volume with manually defined ROI’s using automatic thresholding based on (64), cells were binned as DVG-high or FL-high based on ratio of 3’(-)SeV and 5’(-)SeV.

### Immunoprecipitation

Cells were lysed in NP-40 lysis buffer (Amresco) with proteinase inhibitors (Roche Boehringer Mannheim), RNase OUT (Invitrogen) and EDTA (ThermoFisher). Protein concentration was measured using BCA Protein Assay (ThermoFisher). For anti-NP immunoprecipitation: 500 μg of protein was incubated overnight with primary antibody (anti-SeV NP). Protein G Magnetic Beads (EMD Millipore), 30 μl per sample, were blocked overnight with Salmon Sperm DNA (ThermoFisher) in 5% FBS/PBS. Beads were washed and incubated with lysate/antibody for 4 hours at 4°C while rotating. For anti-NP immunopreciptations, beads were added three times sequentially. Beads were then washed with high salt lysis buffer three times, then low salt lysis buffer once. Samples for RNase treatment were stopped here, samples for other applications were boiled in sample buffer for 10 minutes. For Rab11a-GFP immunoprecipitation: Anti-GFP immunoprecipitation was carried out using anti-GFP mAb magnetic beads. 500 μg lysates were incubated overnight with beads, then washed five times with wash buffer (50 mM Tris-HCl (pH 7.5), 150 mM NaCl, 0.05% NP-40).

### Sample preparation for proteomic analysis

All chemicals used for preparation of nanoflow liquid chromatography-tandem mass spectrometry (nLC–MS/MS) samples were sequencing grade and purchased from Sigma-Aldrich (St. Louis, MO), unless otherwise stated. Immunoprecipitated Rab11a-GFP interacting proteins (Rab11a-GFP IP) were eluted from the magnetic beads by the on-beads tryptic digestion. Briefly, the beads were resuspended in 50 μl of 50 mM triethylammonium bicarbonate, pH 8.5 (TEAB, Thermo Fisher Scientific, Waltham, MA) and proteins were reduced using 10 mM dithiothreitol (DTT) for 1 hour at room temperature and alkylated with 20 mM iodoacetamide (IAA) in the dark for 30 minutes at room temperature. Proteins were digested with trypsin (Promega, Madison, WI) at an enzyme-to-substrate ratio of ~1:50 for 12 hours in a thermomixer, shaking at 900 rpm, at room temperature. After digestion, the supernatant was removed and collected into fresh, labelled tubes. Beads were washed twice with 50 μl of the wash buffer (50 mM TEAB pH 8.5, 5% acetonitrile) and all supernatants were merged. The samples were concentrated to the volume of ~100 μl by lyophilization and acetified with trifluoroacetic acid (TFA) to a final concentration of 0.1%. The tryptic peptides were desalted using Poros Oligo R3 RP (PerSeptive Biosystems, Framingham, MA) P200 columns with C18 3 M plug (3 M Bioanalytical Technologies, St. Paul, MN) prior to nLC-MS/MS analysis.

The whole cell proteome samples were processed using the suspension trap (S-Trap, Protifi, Huntington, NY) (65) mini spin column digestion protocol with minor modifications. Briefly, cells were lysed in 300 μl of lysis buffer (5% SDS, 50 mM TEAB pH 7.55, Halt™ protease and phosphatase inhibitor cocktail (Thermo Fisher Scientific, Waltham, MA)) by vortexing and probe tip sonication at 4 °C. The lysate was clarified by centrifugation at 13,000×*g* for 10 minutes, at 4 °C. Protein concentration was measured by Bradford protein assay (Thermo Fisher Scientific) and ~300 μg of reduced and alkylated proteins was subjected to trypsin digestion following the S-Trap manufacturer’s procedure. The peptide solution was pooled, lyophilized, and desalted prior to nLC–MS/MS.

### Nanoflow Liquid Chromatography Tandem Mass Spectrometry (nLC-MS/MS)

The peptide mixture was separated using a Dionex Ultimate 3000 high-performance liquid chromatography (HPLC) system (Thermo Fisher Scientific) equipped with a two-column setup, consisting of a reversed-phase trap column (Acclaim PepMap100 C18, 5 μm, 100 Å, 300 μm i.d. × 5 mm, Thermo Fisher Scientific) and a reversed-phase analytical column (30 cm, 75 μm i.d. 360 μm o.d., in-house packed with Pur C18AQ 3 μm; Dr Maisch). Loading buffer was 0.1% trifluoroacetic acid (Merck Millipore) in water. Buffer A was 0.1% formic acid, and Buffer B was 80% acetonitrile + 0.1% formic acid. The HPLC was coupled online with an Orbitrap Fusion mass spectrometer (Thermo Fisher Scientific, San Jose, CA). The gradient was 135 min from 2% to 36% buffer B at a flow rate of 300 nl/min for Rab11a-GFP IP samples, and 180 min for whole cell proteome samples. The MS instrument was controlled by the Xcalibur software (Thermo Fisher Scientific). The nanoelectrospray ion source (Thermo Fisher Scientific) was used with a spray voltage of 2.2 kV. The ion transfer tube temperature was 275°C. Data acquisition was performed in the Orbitrap for precursor ions. MS survey scans were obtained for the m/z range of 350-1200 in the Orbitrap with maximum ion injection time of 100 ms, automatic gain control target 5 × 10^5^ and a mass resolution of 120,000. MS/MS was performed with a TopSpeed duty cycle set to 3 s. Dynamic exclusion was set to 4 sec. Charge state enabled was 2-6^+^. Higher Collisional Dissociation (HCD) was set to 30. MS/MS was acquired in the ion trap using the Rapid scan mode, an automatic gain control set to 10,000 and a maximum injection time set to 120 msec.

### Proteins Identification and Quantification

The raw mass spectrometer files were processed for protein identification using the Proteome Discoverer (v2.4, Thermo Fisher Scientific) and the Sequest HT algorithm with a peptide mass tolerance of 10 ppm, fragment m/z tolerance of 0.25 Da, and a false discovery rate (FDR) of 1% for proteins and peptides. All peak lists were searched against the UniProtKB/Swiss-Prot database of Human (January 2020, 20,367 entries) and UniProtKB/TrEMBL Sendai virus (Cantell); February 2020, 8 entries) sequences using the parameters as follows: enzyme, trypsin; maximum missed cleavages, 2; fixed modification, carbamidomethylation (C); variable modifications, oxidation (M), protein N-terminus acetylation. Protein quantifications were log2 transformed and normalized using the median of the distribution for each sample. In order to estimate an enrichment of proteins detected only in mock or infection, missing values were imputed using a distribution of values of 30% width and two standard deviations lower than the average of the distribution of valid values. Statistical analyses were performed on three different biological replicates. The sample size was chosen to provide enough statistical power to apply parametric tests (either homoscedastic or heteroscedastic one-tailed t test, depending on the statistical value of the F-test; heteroscedastic if F-test p value<0.05). The t test was considered as valuable statistical test because binary comparisons were performed, and the number of replicates was limited. No samples were excluded as outliers (this applies to all proteomics analyses described in this manuscript). Proteins with t test p value smaller than 0.05 were considered as significantly altered between the two tested conditions. Data distribution was assumed to be normal, but this was not formally tested.

### Western Blot

Whole cell lysate (10μg) or immunoprecipitation products were denatured by boiling for 10 minutes and run on a 4-12% Gradient Bis-Tris Gel (Bio-Rad) before being transferred to a PVDF membrane (Millipore). Membranes were incubated overnight with primary antibody in 5% milk. Membranes was incubated secondary antibody for an hour in 5% milk and developed using Lumi-light western blot substrate (Roche) to detect HRP. Images were acquired using a ChemDoc biomimager (Bio-Rad) and quantified in ImageJ. Primary antibodies: rabbit anti-Rab11a (Invitrogen); mouse anti-C (clone p96-6, a kind gift from Toru Takimoto), SeV NP, P, and M were detected by chicken anti-SeV polyclonal(Abcam). Secondary antibodies: anti-rabbit HRP-conjugated antibody (Cell Signaling), anti-mouse IgG for IP (Abcam), anti-chicken HRP-conjugated antibody (Invitrogen).

### RNase Treatment

Post-immunoprecipitation unbound fractions or substrate bound to magnetic beads were subjected to treatment with a combination of 1U/mL RNase A, V, and T1 (Invitrogen) for 15 minutes at room temperature. RNase reaction was stopped by adding Trizol LS (Invitrogen) to the sample.

### RNA Extraction and RT-qPCR

Cellular and viral RNA was harvested using Trizol (Invitrogen). RNA was reverse transcribed using SuperScript III First Strand Synthesis System (Invitrogen) with OligoDT for mRNA specific amplification or with primer 5’-GGTGAGGAATCTATACGTTATAC-3’ for viral RNA. qPCR was performed with 1:40 dilution of cDNA, SYBR Green (Life Technologies) and 5 μM forward/reverse primers (Invitrogen) on an Applied Biosystems ViiA 7 Real-Time System. Relative copy numbers per cell were calculated by delta-delta Ct and normalized to average cellular GAPDH expression levels. Primer sequences are: SeV NP (F: 5’ TGCCCTGGAAGATGAGTTAG-3’ R: 5’-GCCTGTTGGTTTGTGGTAAG-3’); SeV P/C (F: 5’ - GGATATCCGAGATCAGGTATTGA-3’ R: 5’-GGCCCGGTGTATATTTTGTTT-3’); SeV M F: (5’- GCCATCCCCTACATCAGGAT-3’ R: 5’- GTAACGACCCGGAGCCGCAT-3’); SeV F (F: 5’-CTCCTGAAGATCTCTAAGGCAT-3’ R: 5’- GGATCCCACGAATCGAGGTA3’); SeV HN F: (5’- GACCAGGAAATAAAGAGTGCA-3’ R: 5’-CGATGTATTGGCATATAGCGT-3’); SeV L (F: 5’- TGGTCAGAGATGCAACGAGA-3’ R: 5’-ACCTTTCAAGGACTGGATGC-3’); gSeV (F: 5’- GACCAGGAAATAAAGAGTGCA-3’, R: 5’-CGATGTATTGGCATATAGCGT-3’), DVG-546 (F: 5’-TCCAAGACTATCTTTATCTATGTCC-3’, R: 5’- GGTGAGGAATCTATACGTTATAC-3’).

## ACKNOWLEDGEMENTS

This work was supported by the U.S. National Institutes of Health National Institute of Allergy and Infectious Diseases (grants R01AI13486, R01AI137062, & R21AI127832 to C.B.L., R01AI125536 to B.L., AI118891 to B.G.A., F31-AI133943 & T32-AI007647-16 to K.A, and AI055940 to T.T.)E.G, was supported by the National Science Foundation Graduate Research Fellowship Program (grant 2016222276). P.A.T. was supported by a Canadian Institutes of Health Research (CIHR) Postdoctoral Fellowship. C.T.H. was supported by the Postdoctoral Research Abroad Program sponsored by Ministry of Science and Technology of Taiwan. The Penn Vet Imaging Core Facility is supported by NIH grant S10 RR027128.

---

**Supplementary Table 1. Time-course monitoring of changes in Rab11a-GFP associated host and viral proteins during Sendai virus infection.** Table includes average of protein abundance detected in three biological replicates for mock, 12hpi and 24hpi infected cells; fold change values obtained for proteins identified at 12 hpi or 24hpi compared to mock infected cells; t-test p-value. ***Protein Accession* -** refers to UniProt database; ***#Peptides* -** highlights the number of razor and unique peptides used for protein quantification; ***# Protein Unique Peptides* -** highlights the number of peptide sequences unique for a given protein*; **Coverage [%]** -* represents the percentage of the protein sequence covered by the peptides identified in the MS run; ***PSMs*** - peptide spectrum matches - show the total number of identified spectra assigned to peptide sequences for the protein, including those redundantly identified; ***CV*** - coefficient of variation; ***Stdev*** - standard deviation. For follow up analysis only proteins identified with minimum 2 peptides *(#Peptides;* razor and/or unique) were selected.

**Supplementary Table 2. Quantitative whole-cell proteome analysis of Sendai virus infected cells.** Table includes average of protein abundance detected in three biological replicates for mock, 12hpi and 24hpi infected cells; fold change values obtained for proteins identified at 12 hpi or 24hpi compared to mock infected cells; t-test p-value. ***Protein Accession* -** refers to UniProt database**; *#Peptides* -** highlights the number of razor and unique peptides used for protein quantification; ***# Protein Unique Peptides* -** highlights the number of peptide sequences unique for a given protein*; **Coverage [%]** -* represents the percentage of the protein sequence covered by the peptides identified in the MS run; ***PSMs*** -peptide spectrum matches - show the total number of identified spectra assigned to peptide sequences for the protein, including those redundantly identified; ***CV*** - coefficient of variation; ***Stdev*** - standard deviation.

## REFERENCES

1. Maykowski P, Smithgall M, Zachariah P, Oberhardt M, Vargas C, Reed C, Demmer RT, Stockwell MS, Saiman L. 2018. Seasonality and clinical impact of human parainfluenza viruses. Influenza Other Respir Viruses 12:706–716.

2. Pawelczyk M, Kowalski ML. 2017. The Role of Human Parainfluenza Virus Infections in the Immunopathology of the Respiratory Tract. Curr Allergy Asthma Rep 17:16.

3. Jain S, Self WH, Wunderink RG, Fakhran S, Balk R, Bramley AM, Reed C, Grijalva CG, Anderson EJ, Courtney DM, Chappell JD, Qi C, Hart EM, Carroll F, Trabue C, Donnelly HK, Williams DJ, Zhu Y, Arnold SR, Ampofo K, Waterer GW, Levine M, Lindstrom S, Winchell JM, Katz JM, Erdman D, Schneider E, Hicks LA, McCullers JA, Pavia AT, Edwards KM, Finelli L, Team CES. 2015. Community-Acquired Pneumonia Requiring Hospitalization among U.S. Adults. N Engl J Med 373:415–27.

4. Pneumonia Etiology Research for Child Health Study G. 2019. Causes of severe pneumonia requiring hospital admission in children without HIV infection from Africa and Asia: the PERCH multi-country case-control study. Lancet 394:757–779.

5. Haywood AM. 2010. Membrane uncoating of intact enveloped viruses. J Virol 84:10946–55.

6. Calain P, Roux L. 1993. The rule of six, a basic feature for efficient replication of Sendai virus defective interfering RNA. J Virol 67:4822–30.

7. Kolakofsky D, Roux L, Garcin D, Ruigrok RWH. 2005. Paramyxovirus mRNA editing, the “rule of six” and error catastrophe: a hypothesis. J Gen Virol 86:1869–1877.

8. Egelman EH, Wu SS, Amrein M, Portner A, Murti G. 1989. The Sendai virus nucleocapsid exists in at least four different helical states. J Virol 63:2233–43.

9. Whelan SP, Barr JN, Wertz GW. 2004. Transcription and replication of nonsegmented negative-strand RNA viruses. Curr Top Microbiol Immunol 283:61–119.

10. Curran J, Marq JB, Kolakofsky D. 1995. An N-terminal domain of the Sendai paramyxovirus P protein acts as a chaperone for the NP protein during the nascent chain assembly step of genome replication. J Virol 69:849–55.

11. Garcin D, Latorre P, Kolakofsky D. 1999. Sendai virus C proteins counteract the interferon-mediated induction of an antiviral state. J Virol 73:6559–65.

12. Irie T, Okamoto I, Yoshida A, Nagai Y, Sakaguchi T. 2014. Sendai virus C proteins regulate viral genome and antigenome synthesis to dictate the negative genome polarity. J Virol 88:690–8.

13. Chang A, Dutch RE. 2012. Paramyxovirus fusion and entry: multiple paths to a common end. Viruses 4:613–36.

14. El Najjar F, Schmitt AP, Dutch RE. 2014. Paramyxovirus glycoprotein incorporation, assembly and budding: a three way dance for infectious particle production. Viruses 6:3019–54.

15. Ke Z, Strauss JD, Hampton CM, Brindley MA, Dillard RS, Leon F, Lamb KM, Plemper RK, Wright ER. 2018. Promotion of virus assembly and organization by the measles virus matrix protein. Nat Commun 9:1736.

16. Tomita Y, Yamashita T, Sato H, Taira H. 1999. Kinetics of interactions of sendai virus envelope glycoproteins, F and HN, with endoplasmic reticulum-resident molecular chaperones, BiP, calnexin, and calreticulin. J Biochem 126:1090–100.

17. Bruce EA, Stuart A, McCaffrey MW, Digard P. 2012. Role of the Rab11 pathway in negative-strand virus assembly. Biochem Soc Trans 40:1409–15.

18. Grant BD, Donaldson JG. 2009. Pathways and mechanisms of endocytic recycling. Nat Rev Mol Cell Biol 10:597–608.

19. Stone R, Hayashi T, Bajimaya S, Hodges E, Takimoto T. 2016. Critical role of Rab11a-mediated recycling endosomes in the assembly of type I parainfluenza viruses. Virology 487:11–8.

20. Nakatsu Y, Ma X, Seki F, Suzuki T, Iwasaki M, Yanagi Y, Komase K, Takeda M. 2013. Intracellular transport of the measles virus ribonucleoprotein complex is mediated by Rab11A-positive recycling endosomes and drives virus release from the apical membrane of polarized epithelial cells. J Virol 87:4683–93.

21. Katoh H, Nakatsu Y, Kubota T, Sakata M, Takeda M, Kidokoro M. 2015. Mumps Virus Is Released from the Apical Surface of Polarized Epithelial Cells, and the Release Is Facilitated by a Rab11-Mediated Transport System. J Virol 89:12026–34.

22. Stricker R, Mottet G, Roux L. 1994. The Sendai virus matrix protein appears to be recruited in the cytoplasm by the viral nucleocapsid to function in viral assembly and budding. J Gen Virol 75 (Pt 5):1031–42.

23. Iwasaki M, Takeda M, Shirogane Y, Nakatsu Y, Nakamura T, Yanagi Y. 2009. The matrix protein of measles virus regulates viral RNA synthesis and assembly by interacting with the nucleocapsid protein. J Virol 83:10374–83.

24. Sanderson CM, McQueen NL, Nayak DP. 1993. Sendai virus assembly: M protein binds to viral glycoproteins in transit through the secretory pathway. J Virol 67:651–63.

25. Hasan MK, Kato A, Muranaka M, Yamaguchi R, Sakai Y, Hatano I, Tashiro M, Nagai Y. 2000. Versatility of the accessory C proteins of Sendai virus: contribution to virus assembly as an additional role. J Virol 74:5619–28.

26. Sakaguchi T, Kato A, Sugahara F, Shimazu Y, Inoue M, Kiyotani K, Nagai Y, Yoshida T. 2005. AIP1/Alix is a binding partner of Sendai virus C protein and facilitates virus budding. J Virol 79:8933–41.

27. Irie T, Nagata N, Yoshida T, Sakaguchi T. 2008. Recruitment of Alix/AIP1 to the plasma membrane by Sendai virus C protein facilitates budding of virus-like particles. Virology 371:108–20.

28. Gosselin-Grenet AS, Marq JB, Abrami L, Garcin D, Roux L. 2007. Sendai virus budding in the course of an infection does not require Alix and VPS4A host factors. Virology 365:101–12.

29. Genoyer E, Lopez CB. 2019. The Impact of Defective Viruses on Infection and Immunity. Annu Rev Virol 6:547–566.

30. Calain P, Roux L. 1988. Generation of measles virus defective interfering particles and their presence in a preparation of attenuated live-virus vaccine. J Virol 62:2859–66.

31. Santak M, Markusic M, Balija ML, Kopac SK, Jug R, Orvell C, Tomac J, Forcic D. 2015. Accumulation of defective interfering viral particles in only a few passages in Vero cells attenuates mumps virus neurovirulence. Microbes Infect 17:228–36.

32. Welch SR, Tilston NL, Lo MK, Whitmer SLM, Harmon JR, Scholte FEM, Spengler JR, Duprex WP, Nichol ST, Spiropoulou CF. 2020. Inhibition of Nipah Virus by Defective Interfering Particles. J Infect Dis doi:10.1093/infdis/jiz564.

33. Kolakofsky D. 1976. Isolation and characterization of Sendai virus DI-RNAs. Cell 8:54755.

34. Tapia K, Kim WK, Sun Y, Mercado-Lopez X, Dunay E, Wise M, Adu M, Lopez CB. 2013. Defective viral genomes arising in vivo provide critical danger signals for the triggering of lung antiviral immunity. PLoS pathogens 9:e1003703.

35. Vignuzzi M, Lopez CB. 2019. Defective viral genomes are key drivers of the virus-host interaction. Nat Microbiol 4:1075–1087.

36. Calain P, Roux L. 1995. Functional characterisation of the genomic and antigenomic promoters of Sendai virus. Virology 212:163–73.

37. Yount JS, Gitlin L, Moran TM, Lopez CB. 2008. MDA5 Participates in the Detection of Paramyxovirus Infection and Is Essential for the Early Activation of Dendritic Cells in Response to Sendai Virus Defective Interfering Particles. J Immunol 180:4910–8.

38. Yount JS, Kraus TA, Horvath CM, Moran TM, Lopez CB. 2006. A novel role for viral-defective interfering particles in enhancing dendritic cell maturation. J Immunol 177:4503–13.

39. Mottet G, Roux L. 1989. Budding efficiency of Sendai virus nucleocapsids: influence of size and ends of the RNA. Virus Res 14:175–87.

40. Xu J, Sun Y, Li Y, Ruthel G, Weiss SR, Raj A, Beiting D, Lopez CB. 2017. Replication defective viral genomes exploit a cellular pro-survival mechanism to establish paramyxovirus persistence. Nat Commun 8:799.

41. Genoyer E, Lopez CB. 2019. Defective Viral Genomes Alter How Sendai Virus Interacts with Cellular Trafficking Machinery, Leading to Heterogeneity in the Production of Viral Particles among Infected Cells. J Virol 93.

42. Tuffereau C, Roux L. 1988. Direct adverse effects of Sendai virus DI particles on virus budding and on M protein fate and stability. Virology 162:417–26.

43. Chambers R, Takimoto T. 2010. Trafficking of Sendai virus nucleocapsids is mediated by intracellular vesicles. PLoS One 5:e10994.

44. Sekellick MJ, Marcus PI. 1980. Viral interference by defective particles of vesicular stomatitis virus measured in individual cells. Virology 104:247–52.

45. McLain L, Armstrong SJ, Dimmock NJ. 1988. One defective interfering particle per cell prevents influenza virus-mediated cytopathology: an efficient assay system. J Gen Virol 69 (Pt 6):1415–9.

46. Wignall-Fleming EB, Hughes DJ, Vattipally S, Modha S, Goodbourn S, Davison AJ, Randall RE. 2019. Analysis of Paramyxovirus Transcription and Replication by High-Throughput Sequencing. J Virol 93.

47. Noton SL, Fearns R. 2015. Initiation and regulation of paramyxovirus transcription and replication. Virology 479-480:545–54.

48. Latorre P, Cadd T, Itoh M, Curran J, Kolakofsky D. 1998. The various Sendai virus C proteins are not functionally equivalent and exert both positive and negative effects on viral RNA accumulation during the course of infection. J Virol 72:5984–93.

49. Utley TJ, Ducharme NA, Varthakavi V, Shepherd BE, Santangelo PJ, Lindquist ME, Goldenring JR, Crowe JE, Jr. 2008. Respiratory syncytial virus uses a Vps4-independent budding mechanism controlled by Rab11-FIP2. Proc Natl Acad Sci U S A 105:10209–14.

50. Bruce EA, Digard P, Stuart AD. 2010. The Rab11 pathway is required for influenza A virus budding and filament formation. J Virol 84:5848–59.

51. Portner A, Murti KG, Morgan EM, Kingsbury DW. 1988. Antibodies against Sendai virus L protein: distribution of the protein in nucleocapsids revealed by immunoelectron microscopy. Virology 163:236–9.

52. Yamada H, Hayata S, Omata-Yamada T, Taira H, Mizumoto K, Iwasaki K. 1990. Association of the Sendai virus C protein with nucleocapsids. Arch Virol 113:245–53.

53. Amorim MJ, Bruce EA, Read EK, Foeglein A, Mahen R, Stuart AD, Digard P. 2011. A Rab11-and microtubule-dependent mechanism for cytoplasmic transport of influenza A virus viral RNA. J Virol 85:4143–56.

54. Bhagwat AR, Le Sage V, Nturibi E, Kulej K, Jones J, Guo M, Tae Kim E, Garcia BA, Weitzman MD, Shroff H, Lakdawala SS. 2020. Quantitative live cell imaging reveals influenza virus manipulation of Rab11A transport through reduced dynein association. Nat Commun 11:23.

55. Curran J, Latorre P, Kolakofsky D. Translational gymnastics on the Sendai virus P/C mRNA, p 351–357. In (ed), Elsevier,

56. Liljeroos L, Huiskonen JT, Ora A, Susi P, Butcher SJ. 2011. Electron cryotomography of measles virus reveals how matrix protein coats the ribonucleocapsid within intact virions. Proc Natl Acad Sci U S A 108:18085–90.

57. Coronel EC, Takimoto T, Murti KG, Varich N, Portner A. 2001. Nucleocapsid incorporation into parainfluenza virus is regulated by specific interaction with matrix protein. J Virol 75:1117–23.

58. Chen W, Feng Y, Chen D, Wandinger-Ness A. 1998. Rab11 is required for trans-golgi network-to-plasma membrane transport and a preferential target for GDP dissociation inhibitor. Mol Biol Cell 9:3241–57.

59. de Castro Martin IF, Fournier G, Sachse M, Pizarro-Cerda J, Risco C, Naffakh N. 2017. Influenza virus genome reaches the plasma membrane via a modified endoplasmic reticulum and Rab11-dependent vesicles. Nat Commun 8:1396.

60. Dimmock NJ, Easton AJ. 2014. Defective Interfering Influenza Virus RNAs: Time To Reevaluate Their Clinical Potential as Broad-Spectrum Antivirals? Journal of virology 88:5217–27.

61. Roux L, Waldvogel FA. 1983. Defective interfering particles of Sendai virus modulate HN expression at the surface of infected BHK cells. Virology 130:91–104.

62. Laske T, Heldt FS, Hoffmann H, Frensing T, Reichl U. 2016. Modeling the intracellular replication of influenza A virus in the presence of defective interfering RNAs. Virus Res 213:90–99.

63. Beaty SM, Park A, Won ST, Hong P, Lyons M, Vigant F, Freiberg AN, tenOever BR, Duprex WP, Lee B. 2017. Efficient and Robust Paramyxoviridae Reverse Genetics Systems. mSphere 2.

64. Costes SV, Daelemans D, Cho EH, Dobbin Z, Pavlakis G, Lockett S. 2004. Automatic and quantitative measurement of protein-protein colocalization in live cells. Biophys J 86:3993–4003.

65. HaileMariam M, Eguez RV, Singh H, Bekele S, Ameni G, Pieper R, Yu Y. 2018. S-Trap, an Ultrafast Sample-Preparation Approach for Shotgun Proteomics. J Proteome Res 17:2917–2924.

